# Matrix prior for data transfer between single cell data types in latent Dirichlet allocation

**DOI:** 10.1101/2022.11.23.517534

**Authors:** Alan Min, Timothy Durham, Louis Gevirtzman, William Stafford Noble

## Abstract

Single cell ATAC-seq (scATAC-seq) enables the mapping of regulatory elements in fine-grained cell types. Despite this advance, analysis of the resulting data is challenging, and large scale scATAC-seq data are difficult to obtain and expensive to generate. This motivates a method to leverage information from previously generated large scale scATAC-seq or scRNA-seq data to guide our analysis of new scATAC-seq datasets. We analyze scATAC-seq data using latent Dirichlet allocation (LDA), a Bayesian algorithm that was developed to model text corpora, summarizing documents as mixtures of topics defined based on the words that distinguish the documents. When applied to scATAC-seq, LDA treats cells as documents and their accessible sites as words, identifying “topics” based on the cell type-specific accessible sites in those cells. Previous work used uniform symmetric priors in LDA, but we hypothesized that nonuniform matrix priors generated from LDA models trained on existing data sets may enable improved detection of cell types in new data sets, especially if they have relatively few cells. In this work, we test this hypothesis in scATAC-seq data from whole *C. elegans* nematodes and SHARE-seq data from mouse skin cells. We show that nonsymmetric matrix priors for LDA improve our ability to capture cell type information from small scATAC-seq datasets.

## 1 Introduction

Single cell genomics has emerged as a powerful method to characterize gene expression (scRNA-seq) and chromatin accessibility (scATAC-seq). The resulting data enables fine-grained identification of cell types. For example, in *Caenorhabditis elegans*, scRNA-seq and scATAC-seq have been used to measure genomewide gene expression levels and chromatin accessibility for the majority of individual cells in the developing embryo and second-stage larval (L2) worms [Durham et al., 2021, Cao et al., 2017, Packer et al., 2019].

Several research groups have found that a Bayesian modeling approach called latent Dirichlet allocation (LDA) is an effective method for distinguishing different cell types in scRNA-seq and scATAC-seq data [Gonzàlez-Blas et al., 2019, Dey et al., 2017]. LDA was developed to model topics in text corpora using counts of words in each document, but when applied to scATAC-seq data, can be used to condense peaks into topics that describe cell types within the data. When applied to scATAC-seq data, the outputs of LDA are a cell-topic matrix, describing the topics assigned to each cell, and a topic-peak matrix, describing how strongly a peak contributes to the definition of each topic. LDA is also well-suited to model single cell genomics data because it expects a matrix of integers as input, and thus can naturally operate on the raw count matrices generated by scATAC-seq or scRNA-seq.

Despite promising results, the challenges posed by scATAC-seq data motivated us to incorporate auxiliary data into the LDA algorithm. Single cell data is still expensive to gather, but there are large compendia of single cell ATAC-seq data available. We aim to use large reference sets of scATAC-seq data (“atlases”) to improve the analysis of smaller datasets through the use of LDA with a nonuniform matrix prior. Specifically, we propose to use previously generated data to create a probabilistic prior for use by the scATAC-seq LDA model. We further investigate the possibility of using scRNA-seq data to transfer information to smaller scATAC-seq datasets. In general, the Bayesian prior methodology provides a principled and computationally lightweight way to incorporate auxiliary data.

We verified the utility of our approach via simulation and then applied the technique to a dataset from *C. elegans* produced using the sci-ATAC-seq assay [Durham et al., 2021] and a dataset from mouse skin cells produced using the SHARE-seq assay [Ma et al., 2020]. We first used simulated data to verify the feasibility of transferring information between two datasets with the same, known, underlying topics; and we show that the nonuniform matrix prior can increase the ability of LDA to identify true underlying topic structure within a given dataset. Next, for the *C. elegans* and mouse skin scATAC-seq data, we split each full data set into a larger “reference” subset and a smaller “target” subset, then applied LDA with a uniform symmetric prior to the reference subset and used the results of that LDA as a nonuniform matrix prior for an LDA model of the target subset. We report that in the mouse skin data, agreement with previously called cell types improved by using auxiliary scATAC-seq data, and that correlation of the output matrices from the “target” LDA with the output matrices from the LDA on the full data set is higher with the nonuniform matrix prior than with the uniform symmetric prior. For the *C. elegans* data, we also found increased correlation of the output matrices between the full data set LDA and the target LDA when using the matrix prior; however, unlike with the SHARE-seq data, we saw no improvement in the agreement with previously called cell types. Finally, we leveraged the paired nature of the scATAC-seq and scRNA-seq data in the mouse skin SHARE-seq dataset to attempt to transfer information across single cell assays. We used the output from an LDA on the scRNA-seq data as a matrix prior for an LDA on the scATAC-seq data. In this case, we did not see a clear improvement when using the matrix prior, but the cross-assay matrix prior might still be improved with further hyperparameter tuning.

## 2 Approach

### 2.1 Background: latent Dirichlet allocation

LDA was originally developed to model text documents as a mixture of topics. For a fixed vocabulary, a document is described as the number of occurrences of each word in the vocabulary. In our case, instead of words, we are modeling scATAC-seq peaks or scRNA-seq genes. We assume that each cell is generated from a mixture of topics, and that each topic has a distribution of peaks or genes from the vocabulary associated with it. The parameters we hope to estimate are the distribution of topics for each cell and the distribution of peaks or genes for each topic, called the “cell-topic matrix” and the “topic-peak matrix” or “topic-gene matrix,” respectively. A generative model is assumed for each of the cells, described below, following [Blei et al., 2003].

Let *N* be the number of sequencing reads or unique molecular identifiers (UMIs) in a cell. Let *T* be the number of topics. Let *V* be the number of peaks or genes in the vocabulary. Let *U* be the number of cells. Let the vector **w** = (*w*_1_,…, *w*_*N*_) be a cell, where each *w*_*i*_ is the number of reads or UMIs associated with a particular peak or gene from the vocabulary. Let ***θ*** = (*θ*_1_,…, *θ*_*T*_) be the topic distribution of a given cell, where 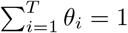. Let *ϕ*_*t*_ = (*ϕ*_*t*1_,…, *ϕ*_*tV*_) be the peak or gene distribution for topic *t* such that 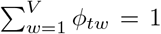, where *ϕ*_*tw*_ indicates the probability of observing a read or UMI from the peak or gene *w* given that its topic was *t*. Let ***α*** be the “basis vector” of length *T* such that ∑_*i*_ *α*_*i*_ = 1 and *α*_*i*_ > 0 for each *i*, which parameterizes the Dirichlet prior distribution over the cell-topic matrix. Similarly, let ***β*** be a basis vector of length *V* such that ∑_*j*_ *β*_*j*_ = 1 and *β*_*j*_ > 0 for each *j*, which parameterizes the Dirichlet distribution over the topic-peak or topic-gene matrix. Let *c*_*α*_ be the “concentration parameter” for ***α***, and *c*_*β*_ be the concentration parameter for ***β***. Let *ξ* be the average number of peaks or genes in a cell.

We note that ***α*** is a prior that describes the frequency at which topics are observed, and ***β*** is a prior that describes the frequency at which peaks or genes are observed, i.e. a large value of *α*_*i*_ or *β*_*i*_ indicates that the *i*th topic or *i*th peak or gene is more common, respectively.

Blei et al. [2003] first describe a variational Bayes approach to optimizing the posterior distribution of the topic assignments for each peak or gene. Following Gonzàlez-Blas et al. [2019] and Darling [2011], however, we use a Gibbs sampling algorithm for LDA for scATAC-seq data (Algorithm 1).

#### Algorithm 1

LDA Gibbs sampling algorithm to estimate topic assignments for each word, by calculating *p*(*z*_*ij*_ = *k* | ***z***^(− *i*)^, ***w***), the probability of topic *z*_*ij*_ being *k* given all other topic assignments and all documents

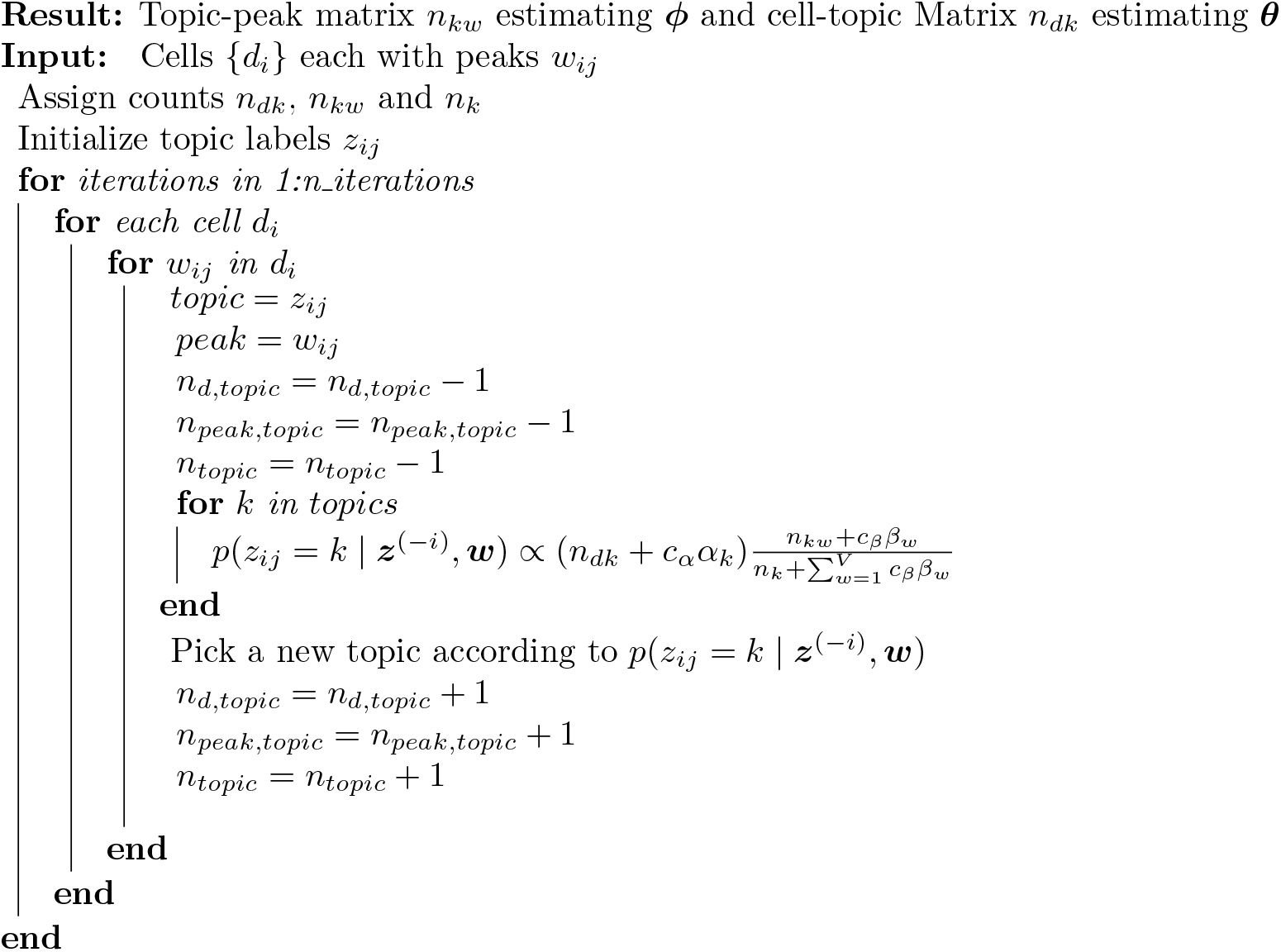

In Gonzàlez-Blas et al. [2019], the last Gibbs sampling iteration is chosen to be the output of the topic-peak and cell-topic matrices. We modify this approach slightly by calculating the posterior likelihood of each of the iterations and choosing the iteration with the highest likelihood. This is the maximum a posteriori (MAP) estimate.

### 2.2 Data transfer using a matrix prior

To achieve our goal of leveraging large scale data to improve inference in small datasets, we extend Algorithm 1 to accept a matrix prior ***B*** to replace the vector prior ***β*** for the topic-peak or topic-gene distribution. Unlike the vector prior ***β***, which specifies the same prior distribution over vocabulary elements regardless of topic, the matrix prior ***B*** can specify a different prior distribution for each topic. The matrix prior ***B*** has elements *B*_*tw*_ such that for each *t*, ∑_*w*_ *B*_*tw*_ = 1, corresponding to the probability of observing peak or gene *w* in topic *t*. It has corresponding concentration parameter *c*_*B*_. We modify the distribution of topics in Algorithm 1, *p*(*z*_*ij*_ = *k* | ***z***^(*-i*)^, ***w***), the probability of topic *z*_*ij*_ being *k* given all other topic assignments and all documents, in the algorithm by instead writing that

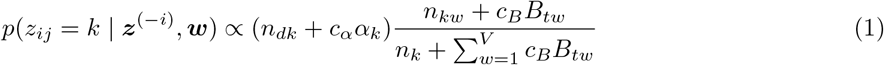

An interpretation of *c*_*B*_*B*_*tw*_ is that we are adding pseudocounts of the peak or gene *w* to the topic *t*. This gives us finer control over the prior distribution of the topic-peak or topic-gene matrix. A key feature of using a matrix prior is that we can use the inferred *ϕ* from one LDA model as the prior for another model with the same vocabulary and similar topics, thereby transferring information from one model to another. We note that this proposed method is similar to a method first proposed in [Wood et al., 2017], but the exact form of the prior is different due to the hyperparameter *c*_*B*_.

We call the dataset that we use to generate the matrix prior the “reference dataset” and the dataset to which we apply the prior the “target dataset.” In order to transfer information from the reference to the target, we first run the LDA algorithm on the reference, where both ***α*** and ***β*** are set to be the “uniform priors” described in Algorithm 1; in other words, *α*_*i*_ = 1*/T* and *β*_*j*_ = 1*/V*. This outputs an estimate of the topic-peak or topic-gene matrix, 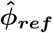, that contains information about the distribution of peaks or genes in each topic. We set the prior 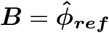, with a concentration parameter *c*_*B*_. As the concentration parameter increases, the LDA algorithm upweights 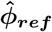.

## 3 Methods

### 3.1 Data

#### 3.1.1 Simulated data

To evaluate the performance of the matrix prior methodology, we first simulated scATAC-seq data and compared inferred topics to the true generative topics that we used to create the synthetic data according to the LDA generative process (Section 2.1). We fixed the true topic-gene distribution across all simulated datasets, with peak or gene distributions for each topic simulated with a Dirichlet distribution with parameters *c*_*β*_ = 0.1, *β*_*j*_ = 1*/V*, resulting in *V*-element vectors. We simulated a new cell-topic matrix for each new dataset, also using a Dirichlet distribution, with parameters *c*_*α*_ = 0.3, *α*_*i*_ = 1*/T* resulting in a *T*-element vector. We set the number of peaks/genes *V* = 8000, the number of topics *T* = 30, and the number of reads/UMIs per cell to be on average *ξ* = 4000.

In the *true matrix simulation*, the goal was to evaluate the matrix prior when the true topic-gene matrix was used as the matrix prior in LDA of simulted data. We simulated target datasets with 1000, 2000, 4000, and 8000 cells, all with the same topic-gene matrix. For each dataset size, we simulated five datasets. In the *inferred matrix simulation*, instead of providing the true topic-gene matrix, we inferred the topic-gene matrix by performing an LDA of simulated reference data using a uniform symmetric prior, then provided that inferred matrix as the prior to LDA of simulated target data sets. We generated four synthetic reference datasets: one for each of 1000, 2000, 4000, or 8000 cells, and we simulated four target datasets of 1000 cells. We performed LDA with a uniform symmetric prior on each reference dataset, and used each resulting topic-gene output matrix as a prior for LDA of each target dataset.

#### 3.1.2 *C. elegans* scATAC-seq data

Recently published *C. elegans* scATAC-seq data demonstrated increased resolution of cell types compared to bulk ATAC-seq data [Durham et al., 2021]. The cells were collected from animals in larval stage 2, and the authors used LDA followed by clustering to identify cell types, which are the labels we use here. The dataset has 30,764 cells and 13,734 peaks. We split the cells uniformly at random into a “reference dataset” of 27,764 cells and a “target dataset” of 3,000 cells to investigate the performance of the matrix prior.

#### 3.1.3 SHARE-seq mouse skin data

SHARE-seq is a co-assay that generates both scATAC-seq and scRNA-seq data from the same single cells simultaneously [Ma et al., 2020]. For our analysis we selected the mouse skin data set from the original publication, which has 34,774 cells from 22 cell types. The scATAC-seq data for these cells consist of chromatin accessibility across 344,592 peaks, while the scRNA-seq data contain the expression measurements of 22,813 genes. First, we repeated our experiments from the *C. elegans* dataset, testing the effectiveness of the matrix prior for transferring information from a larger reference dataset to a smaller target dataset. Second, we leveraged the co-assay data, in which we know the ground-truth pairing of scATAC-seq and scRNA-seq measurements for each cell, to investigate whether we could use the matrix prior to transfer information between scRNA-seq and scATAC-seq datasets.

### 3.2 LDA analysis

#### 3.2.1 LDA analysis of simulated data

For our analysis of simulated data, we used the implementation of the LDA algorithm that employs the Gibbs sampling scheme described in Algorithm 1 [Durham et al., 2021]. We set the number of Gibbs sampling iterations to be 1000. The iteration with the highest posterior probability was used to infer the cell-topic and topic-gene matrices.

In the true matrix simulation, we used the true generative topic-gene matrix as the matrix prior for the target data LDA. We compared the quality of the inferred cell-topic and topic-gene matrices both using the matrix prior and using the uniform prior. For the uniform prior, we supplied LDA with the parameters *C*_*α*_ = 0.3, and *c*_*β*_ = 0.1, matching the simulation Dirichlet parameters. When we used the matrix prior, we supplied LDA with the concentration parameter *c*_*α*_ = 0.3, which matched the simulation Dirichlet parameter, and we set the matrix prior concentration parameter to *c*_*B*_ = 1000.

In the inferred matrix simulation, we used LDA with a uniform prior to infer a topic-gene matrix 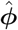 using each of the four reference datasets with between 1000 and 8000 cells. We trained these uniform prior LDA models with *c*_*α*_ = 0.3, and *c*_*β*_ = 0.1. We then used each resulting 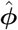 as the matrix prior for each of the four target datasets. We again compared using the matrix prior to using a uniform prior. We used the same settings for the matrix prior LDA models as in the true matrix simulation.

#### 3.2.2 LDA analysis with experimental data: transfer from scATAC-seq to scATAC-seq

To assess whether using a matrix prior would improve LDA performance compared to a uniform prior on small, sparse data, we first trained an LDA model with a uniform prior on all available cells to compare to using our prior on a small dataset. We will refer to this model as the “joint model”. Next, we split the data into a larger “reference set” of 31,774 cells and a smaller “target set” of 3000 cells, and trained a uniform prior LDA model on each of these sets. The output from the target data when the uniform prior was used set the floor for the expected performance of our prior. Last, we used the gene-topic probabilities from the reference model as a matrix prior for a new LDA model of the target set of cells, and we compared the results of this model with those of the uniform models to evaluate the effectiveness of the matrix prior.

#### 3.2.3 Hyperparameter search

We conducted a hyperparameter search to inform our selection of values for the number of topics to use, *T*, and the two concentration parameters *c*_*α*_ and *c*_*B*_. We used a grid-search strategy, testing seven values for *T* (2, 3, 4, 5, 10, 15, and 20), four values for *c*_*α*_ (0.03, 0.3, 3.0, and 30.0), and eight values for *c*_*B*_ (10, 50, 75, 150, 250, 1000, 2000, and 4000); and we employed a likelihood-based measure, perplexity, as our evaluation metric, following Wallach et al. [2009]. Perplexity is the negative exponent of the likelihood, and is calculated for each cell as 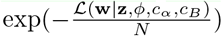, where *N* is the number of reads in a cell. A lower value of perplexity is better. To calculate the likelihood required for the perplexity measure, we use the Chib-style estimation procedure [Chib, 1995], which is a method using a Markov chain to evaluate ℒ(**w** | **z**, *ϕ, c*_*α*_, *c*_*B*_).

Due to our two-tiered method, in which we train a uniform prior LDA on a reference subset of the data and then use the output as a matrix prior for an LDA on the target subset of the data, we also required two corresponding tiers for our hyperparameter search. Thus for the first tier, we generated a set of matrix priors by training, for each value of *T* in our grid search, a uniform prior LDA model on the reference data set. All of these models used *c*_*α*_ = 3 and *c*_*β*_ = 800, which we chose based on the hyperparameter values reported in Durham et al. [2021]; note that we lowered *c*_*β*_ compared to the published value of 2000 to allow the data to have a greater role in determining the topic-gene matrices we would use as matrix priors.

In the second tier of the hyperparameter search, we conducted the full grid search on the target data set with a range of values for *T, c*_*α*_, and *c*_*B*_. We split the target data into ten different training/test splits of 2700 and 300 cells, and then trained each of the ten splits on each of the hyperparameter combinations, with the LDA using the matrix prior from tier 1 that corresponded to each value of *T*. Then, we evaluated the performance of each hyperparameter combination by computing the perplexity on the held out test set cells.

We found that the optimal number of topics was 10, that the perplexity was relatively insensitive to the choice of *c*_*α*_, and that perplexity dropped with increasing values of *c*_*B*_. Following Durham et al. [2021], we increased the number of topics to add some flexibility to the model, and set *T* to be 15 in our experiments. The results of the hyperparameter search and further comments are in Figures S1 and S2. UMAP plots of the results of LDA for different numbers of topics are shown in Figure S3.

We conducted the hyperparameter search for our LDA modeling of the SHARE-seq mouse skin data in similar fashion to the hyperparameter search for the *C. elegans* data. However, the SHARE-seq scATAC-seq dataset contains about ten times more reads per cell than the *C. elegans* dataset, leading to much longer LDA training times per cell and higher memory usage. To make the hyperparameter search more efficient, we limited the SHARE-seq analysis to the 20,000 peaks with the highest variance amongst cells, and randomly downsampled the dataset to 7,000 cells. We split the 7000 cells into 6300 cells for the reference dataset, and 700 cells for the target dataset. We then made ten train/test splits of the target data set by sampling 630 training cells and using the remaining 70 held-out test cells for the Chib method. As with the *C. elegans* data, our hyperparameter search results show that perplexity was not very sensitive to the *c*_*α*_ parameter, and that perplexity decreased as the value of the concentration parameter *c*_*B*_ increases, suggesting that the matrix prior was able to improve the quality of the our inference (Figure S2). However, surprisingly, our hyperparameter search achieved the lowest perplexity with just *T* = 2 topics. We conjectured that this may be because of the low number of peaks included in the analysis, the low number of cells, or the peaks with highest variance may not be able to distinguish the cells. It is also possible that without a normalization for the mean accessibility, we might not select for peaks with the most meaningful differences in accessibility. We hence opted to instead continue using 15 topics as in the *C. elegans* case. UMAP plots of the results of LDA for different numbers of topics are shown in Figure S4, and we found that 10-15 topics visually had good separation of the previously called cell types.

#### 3.2.4 LDA analysis with experimental data: transfer from scRNA-seq to scATAC-seq

To use scRNA-seq data to generate a matrix prior for the LDA of the scATAC-seq data, our method requires that the scATAC-seq and scRNA-seq data share the same vocabulary. Hence, we translated the scATAC-seq data from a vocabulary of peaks to one of genes by counting the number of ATAC-seq cut sites per cell that overlapped each gene and its promoter (defined as the 2kb region immediately 5’ of the transcription start site). We defined cut sites as reported in [Durham et al., 2021]; briefly, they are 60 bp regions centered on the mapping locations of the 5’ ends of the paired end reads (which define the extent of the original DNA fragment that was cut out of the genome). We investigated whether this translated scATAC-seq data retained similar information to the native scATAC-seq data by qualitatively comparing UMAP plots of the LDA output of the raw peaks versus the summed cut sites and saw that the plots were qualitatively similar (Figure S5). Furthermore, we investigated the similarity between the scRNA-seq data and the translated scATAC-seq data by testing the correlation between the two data sets based on the number of counts per cell and counts per gene (Figure S6). We found that there was a moderate amount of correlation between the scRNA-seq data and the scATAC-seq data, suggesting that generating a matrix prior from scRNA-seq data and applying it to an LDA of a translated scATAC-seq data set may be useful. See Supplementary note 1 for results and discussion of our work in this direction.

### 3.3 Evaluation

#### 3.3.1 Evaluation of LDA in simulated data

We used the mean squared error (MSE) to evaluate our inferred cell-topic matrix 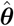 and topic-gene matrix 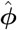. Then, we calculated the MSEs for the cell-topic matrix and topic-gene matrix using

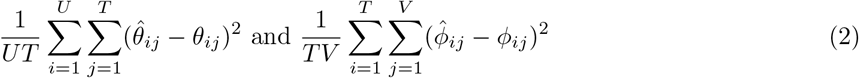

where *U* is the number of cells, and *T* is the number of topics.

In addition to MSE, we calculated Pearson’s *r* and Spearman’s *r* after flattening the cell-topic and topic-gene matrices.

One complication in simulation is that the order of topics inferred by LDA will not necessarily match the order of topics in the simulated ground truth; i.e. topic 1 from the output of LDA is not necessarily semantically the same as topic 1 in the simulated true cell-topic matrix. Therefore, prior to calculating any performance measure, we must match topics. We do this by using a greedy approach on the topic-peak or topic-gene matrix, considering each pair of topics in sorted order by Euclidean distance and allowing only one-to-one matches. For each topic, we match it to the true topic that is closest in MSE.

We used this greedy topic matching algorithm when evaluating the uniform prior LDA models in both the true matrix simulation and the inferred matrix simulation, and when evaluating the matrix prior LDA model for the inferred matrix simulation. We did not need to match topics for the matrix prior LDA in the true matrix simulation, because providing the true topic-gene distributions as the prior already imparts the proper topic semantics to the target LDA, making the inferred topic-gene matrix and the true topic-gene matrix directly comparable.

#### 3.3.2 Evaluation of LDA in experimental data

Unlike our simulation experiments, in which we know the true underlying topic distributions, we do not have a ground truth to compare against for our analyses of datasets derived from biological experiments. Instead, to evaluate the performance of our LDA models, we compared the output of each LDA on a target subset to the LDA output when training on the full data set (i.e. the “joint model”), with the interpretation that if our matrix prior allows the smaller target data set LDA to infer similar topics to the joint model, then our matrix prior has succeeded. To make this comparison, we selected just the rows in the cell-topic matrix of the joint model that corresponded to the target dataset cells, we applied our greedy topic matching algorithm to account for any topic-switching that occurred in the LDA of the reference data set compared to the joint model, and then we evaluated the similarity between the outputs of the matrix prior LDA and the joint model using Pearson’s correlation, Spearman’s correlation, and MSE. We focus on Pearson correlation in the main text but report other metrics in the supplementary Material.

In addition to our quantitative evaluation of the matrix prior approach, we qualitatively evaluated its performance by visualizing the cell-topic matrix using UMAP [McInnes et al., 2018]. For the *C. elegans* data, we took the cell-topic matrix LDA output from one of the splits of the target training set, computed a two dimensional UMAP embedding, and represented the embedding as a scatter plot in which we colored the cells based on their published cell type labels [Durham et al., 2021]. We quantitatively measured how well the inferred cell-topic matrices could separate cells of different types (both at the level of broader cell types and more specific ones, such as neuron subtypes) by using the silhouette coefficient. The silhouette coefficient is a distance-based measure of cluster cohesiveness that has previously been used as a performance metric when comparing the performance of different single cell analysis methods Luecken et al. [2022], and it is computed using Equation 3. For a cell with index *i*, and *C*_*i*_ the set of indices of cells with the same cell type label as cell 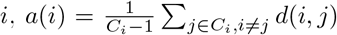 is the average Euclidean distance to all other cells of the same cell type. We similarly define 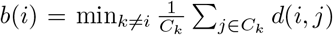 to be the lowest average Euclidean distance to a different cell type from cell *i*.

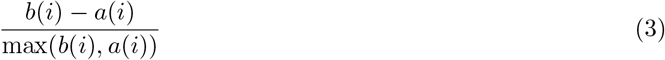

The resulting silhouette coefficent is bounded between −1 and 1. Values close to 1 indicate that a point is closer to points in the same cluster than to points in other clusters, and a high silhouette coefficient indicates that the LDA topics reflect the published cell types well.

### 3.4 Data availability

The code to run LDA with a matrix prior is available at https://github.com/gevirl/LDA using the -betaFile option. There are no primary data in the paper.

## 4 Results

### 4.1 Matrix prior improves topic inference in simulation

We began testing the matrix prior by running LDA on simulated data derived from known cell-topic and topic-gene matrices. In the true matrix simulation, we tested the hypothesis that an LDA given the true topic-gene matrix as a prior would yield results more similar (by MSE) to the ground truth than the uniform prior LDA would. We also tested the hypothesis that increasing the number of target cells would reduce the performance advantage of the matrix prior over the uniform prior. In the inferred matrix simulation, we tried using LDA output on a reference data set as our matrix prior for the target LDA and tested the effect of varying the reference data set size on the performance of the target LDA.

In the true matrix simulation, we found that using the matrix prior instead of the uniform prior led to more accurate inference of the topic-gene and cell-topic matrices (Figure 1a). We kept the weight of the matrix prior, *c*_*B*_, constant and varied the number of cells in the target dataset to understand effects of target dataset size. The MSE was higher when we used a uniform prior than when we used the matrix prior regardless of target dataset size (we tested 1000, 2000, 4000, and 8000 target cells), although the difference in performance decreased as the cell number increased, suggesting that with more cells the data began to overwhelm the prior. To better understand how the matrix prior improves the LDA results, we analyzed some representative uniform prior LDA results in more detail. When the number of cells was low, the uniform prior LDA underestimated the weights of high probability topics, and even at 8000 cells, it incorrectly predicted topic assignments in some cells (Figure S7, top row). On the other hand, for the matrix prior LDA, the inferred and true cell-topic matrices agreed (Figure S7, bottom row). We found similar results for the topic-gene matrices (Figure S8).

**Figure 1:**
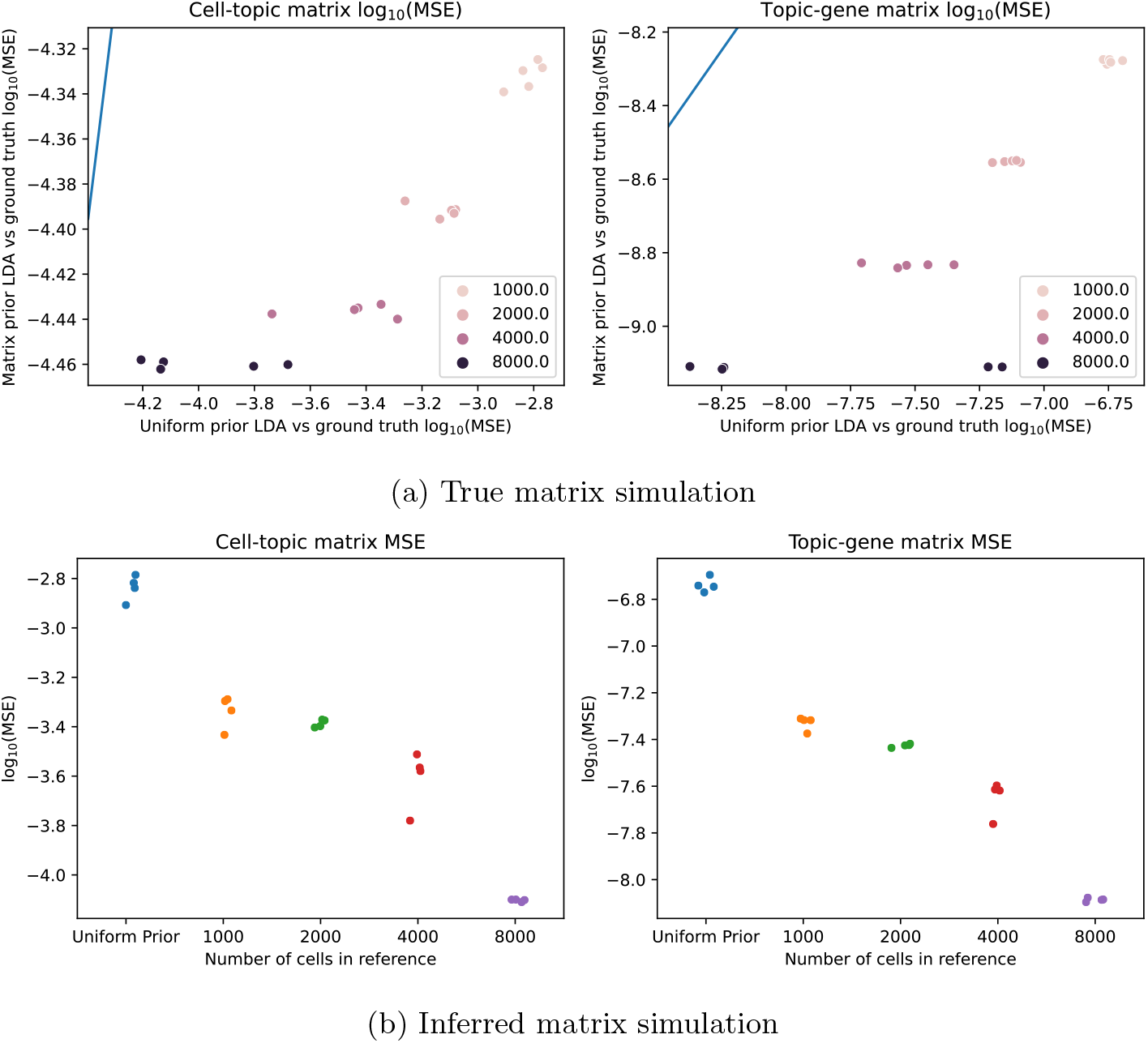
Simulation experiments show that the matrix prior improves the concordance between inferred topics and the ground truth compared to the uniform prior. Experiments from the true matrix simulation and the inferred matrix simulation are shown here for different numbers of cells in the target dataset (different colors). (a) MSE from the ground truth to the LDA with a ground truth matrix prior (y-axis) is plotted against the MSE from the ground truth to the uniform symmetric prior LDA (x-axis), for both the cell-topic matrix (left) and the topic-gene matrix (right). The blue line is the line *y* = *x*. Each point represents one independently simulated dataset, with a unique true cell-topic matrix and topic-gene matrix. (b) MSE to the ground truth for the LDA with a matrix prior inferred from a simulated reference dataset is shown for different reference data set sizes. MSE is plotted for both the cell-topic matrices (left) and the topic-gene matrices (right).

In the inferred matrix simulation, we found that the matrix prior improved the inference of the cell-topic and the topic-gene matrices compared to the uniform prior, as measured by MSE against the ground truth (Figure 1b). We also trained a series of LDA models on a 1000 cell target data set, each with a matrix prior generated from a uniform LDA trained on a reference dataset with 1000, 2000, 4000, or 8000 cells. As the number of reference cells increased, the MSE continually improved (x-axis), and we note that even the matrix prior generated from a reference dataset of only 1000 cells improved LDA performance compared to a uniform prior (Figure 1b). In representative simulated datasets, the cell-topic and topic-gene matrices more closely followed the *y* = *x* line as the number of reference cells increased (Figure S9). These two simulations suggest that under ideal conditions, the matrix prior method is able to improve the quality of inferred topics in LDA.

### 4.2 Whole worm scATAC-seq prior improves concordance with joint model

We used the *C. elegans* data to validate the ability of our matrix prior to improve LDA inference on real data. We randomly split the 34,764 cells into a 3,000 cell target dataset and a 27,764 cell reference dataset, and used the split data to train a matrix prior LDA on the target dataset. Then, we trained a separate uniform prior LDA on the full *C. elegans* dataset (the “joint model”), and compared the inferred topics between the matrix prior LDA and the joint model. We also trained a uniform prior LDA on the target dataset, and evaluated whether the matrix prior LDA results were more similar to the the joint model than the uniform prior LDA results.

We found that the Pearson correlation between the target LDA and the joint model was higher when we used the matrix prior than when we used the uniform prior (Figure 2, left). The correlation increased with increasing *c*_*B*_, but eventually reached a saturation point. We also found that the matrix prior outperformed the uniform prior in Spearman correlation and MSE (Figure S10). When plotting the values of the cell-topic and topic-gene matrices, the points near the *y* = *x* line for the cell-topic matrix tightened as the concentration parameter *c*_*B*_ increased (Figure S14). We also additionally note that in our hyperparameter search, we found evidence that the use of the matrix prior improved the quality of the topic-gene matrix, since the perplexity value in the held out test set improved as the matrix prior concentration parameter increased (Figure S1)

**Figure 2:**
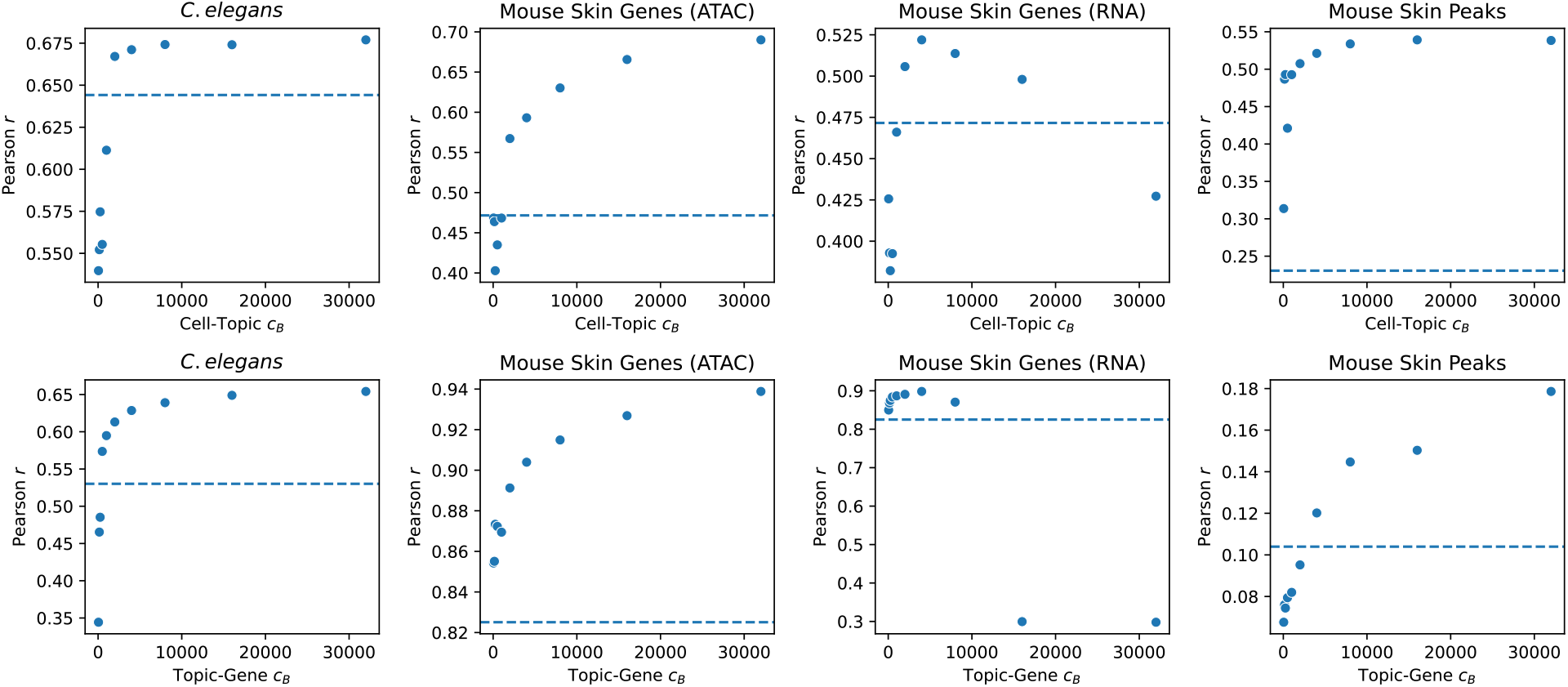
The Pearson correlation between LDA results on the target dataset for the matrix prior LDA and the full dataset uniform symmetric LDA (the “joint model”) increases as *c*_*B*_ increases. Pearson *r* values are plotted as a function of *c*_*B*_ for the cell-topic (top row) and topic-gene (bottom row) matrices for LDA experiments on four different datasets: *C*.*elegans* scATAC-seq data (first column), SHARE-seq mouse skin scATAC-seq data with the peak vocabulary translated to genes (second column), SHARE-seq mouse skin scRNA-seq data (third column), and SHARE-seq mouse skin scATAC-seq data using the peak vocabulary (fourth column). The dotted horizontal lines indicate the correlation between the uniform prior LDA and the joint model.

### 4.3 SHARE-seq scATAC-seq matrix prior improves concordance with the joint model

We used data from mouse skin cells analyzed using the SHARE-seq assay [Ma et al., 2020] to further validate the ability of a matrix prior to improve inference in LDA. We split the 34,774 cells into a 31,774 cell reference dataset and a 3000 cell target dataset, and the same analyses were applied as in the *C. elegans* data (Section 4.2).

We found that, as with the *C. elegans* data, the matrix prior improved our inference in the SHARE-seq mouse skin data. For both the cell-topic and topic-gene matrices, the Pearson correlation of the matrix prior LDA vs the joint model improved as *c*_*B*_ increased (Figure 2, second and fourth columns) and was consistently better than when using the uniform prior (dotted blue line). We saw similar results when using Spearman correlation and MSE as our performance measures (Figures S11, S13). We additionally note that like in the *C. elegans case*, we found evidence that the use of the matrix prior improved the quality of the topic-gene matrix in the hyperparameter search, since the perplexity value in the held out test set improved as the matrix prior concentration parameter increased (Figure S2)

We note that increasing the value of *c*_*B*_ more consistently improved the correspondence between the target LDA and the joint model for the topic-gene matrix than the cell-topic matrix. This difference is most likely due to the fact that the matrix prior is specified as a prior on the topic-gene distribution, and thus only influences the cell-topic distribution indirectly through the training of the LDA model.

In addition to serving as another dataset to validate our scATAC-seq prior, the SHARE-seq co-assay data allowed us to evaluate whether a matrix prior generated from one data modality, scRNA-seq, could improve LDA performance on another data modality, scATAC-seq. Because the SHARE-seq scATAC-seq and scRNA-seq data were generated from the same cells, we were able to directly assess the agreement between the scRNA-seq LDA and the scATAC-seq LDA, with and without a matrix prior derived from the scRNA-seq data (Section 3.2.4). See Supplementary Note 1, where we report that the scRNA-seq prior was able to improve inference for moderate values of *c*_*B*_ but worsened inference for larger values.

### 4.4 Mouse skin cell types are more clearly separated with the use of the matrix prior

We next aimed to assess whether using a matrix prior would improve the ability of LDA to distinguish the cell types in a small subset the SHARE-seq mouse skin data set [Ma et al., 2020]. We split the data into a reference data set and a target data set (see Methods), and then we analyzed the target data set using both a uniform prior and a matrix prior derived from LDA on the reference data set. After applying UMAP to our two models to reduce the 15-dimensional topic space into a two-dimensional UMAP space, we qualitatively observed that the matrix prior LDA resulted in cell clusters that better agreed with the published cell type labels than the uniform prior LDA, and this improvement became more marked as we increased the weight of the prior, *c*_*B*_ (Figure 3). We also used the silhouette score to quantitatively measure how well the cells clustered by their cell type labels (Figure S18). We found that as we trained matrix prior models with increasing values of *c*_*B*_, the silhouette score also increased, indicating that the resulting cell clusters better agreed with the cell type labels. The overall silhouette values ranged from −0.113 using the uniform prior to −0.069 with the highest value of *c*_*B*_.

**Figure 3:**
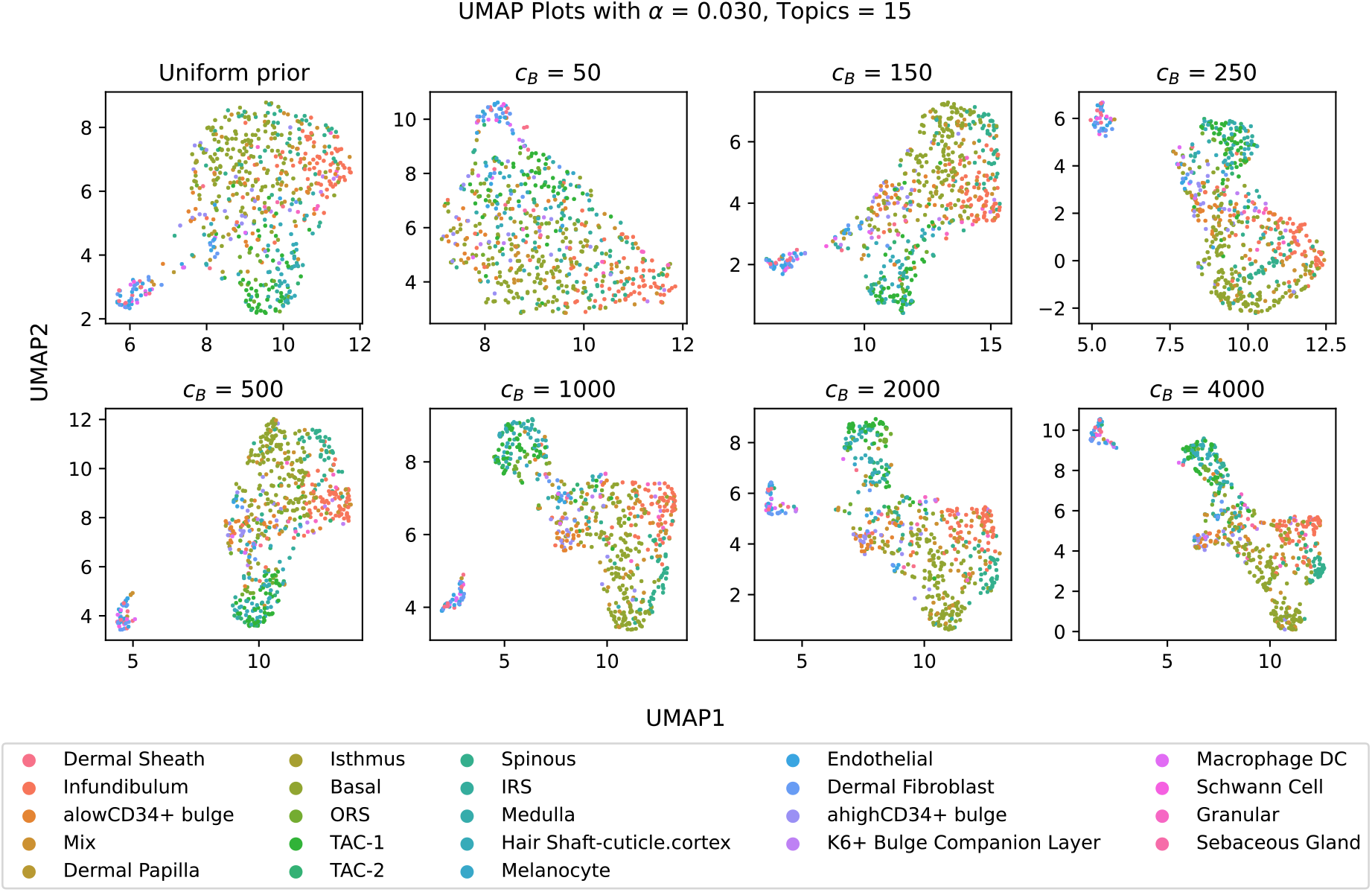
Increasing the weight of the matrix prior shows a qualitative improvement in the ability of the target dataset LDA to discriminate among cell types. UMAP embeddings of the cell-topic matrices from SHARE-seq mouse skin scATAC-seq data using the peak vocabulary are shown for the uniform prior LDA (top left) and matrix prior LDA models trained with different values of *c*_*B*_. Scatter points representing cells are colored by their published cell type annotations.

Note that the silhouette scores are negative, which indicates a high level of cell type overlap in the LDA embedding. A possible explanation for this is that there are many cell types in the SHARE-seq data, some of which are very similar to each other [Ma et al., 2020], and our small target data set (in which we subsampled both cells and peaks) perhaps provides limited power to distinguish among them (Table S1). To investigate further, we repeated the silhouette score analysis with only cells from the top 5 most common cell types, which again showed increasing silhouette scores as *c*_*B*_ increases, but with positive values for the silhouette scores (Figure S19).

We also conducted a similar experiment in *C. elegans data* (Supplementary Note 2); however, we did not see a similar increase in the silhouette coefficient. A possible reason for this is that even in the case of the uniform prior, LDA created separation in the *C. elegans* cell types, and hence no further improvement was possible by using the matrix prior. This may be because the target data set for *C. elegans* has 3000 cells and used all of the peaks, whereas the SHARE-seq data set target data set only included 630 cells and used a subset of the peaks.

## 5 Discussion

We have shown through both simulation study and through analysis of real data that the matrix prior we propose is able to capture information from a larger reference dataset and impart the semantics of the topic-gene or topic-peak matrix onto a smaller target dataset. In our simulation studies, we found that when the true topic-gene and cell-topic matrices were known, we were able to recover those matrices both by directly inputting the truth as the prior and by inferring the topic-gene matrix from a reference data set (Figure 1). These simulations demonstrated that in ideal conditions, the matrix prior can greatly improve performance of LDA. In our real data examples, we examined *C. elegans* and mouse skin cell data. We saw promising results when we used scATAC-seq data to generate a prior for analyzing a target scATAC-seq dataset. We found that LDA results on a small target dataset were more concordant with a model of the full data set when we used a matrix prior derived from a larger reference dataset than when we used a uniform prior (Figure 2). Furthermore, in the case of the mouse skin data, we showed qualitative and quantitative improvements in cell type discrimination for the matrix prior LDA compared to the uniform prior LDA (Figure 4).

**Figure 4:**
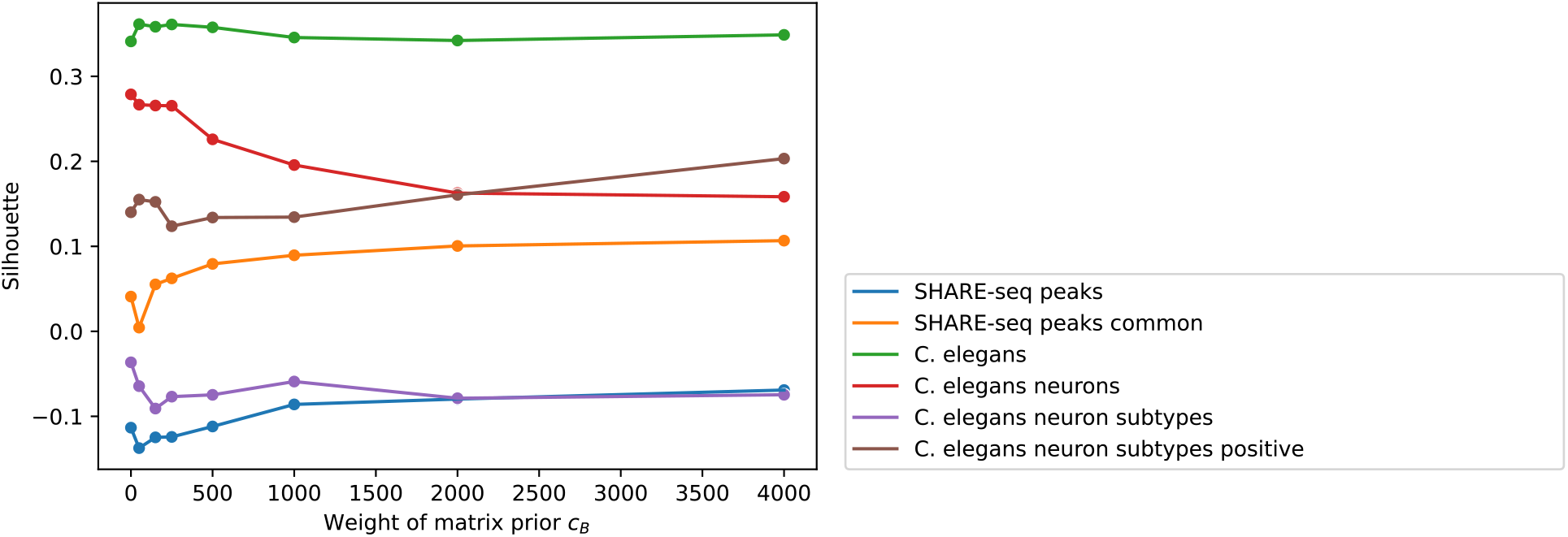
Silhouette values demonstrate quantitative improvement of correspondence with published cell type annotations. Average silhouette values for the published cell type annotations (y-axis) are plotted against increasing values of *c*_*B*_ (x-axis) for six different datasets (colors). The line labeled “SHARE-seq peaks” shows the average silhouette values for the SHARE-seq scATAC-seq data using the peaks vocabulary. “SHARE-seq peaks common” shows the average silhouette values for the SHARE-seq scATAC-seq peaks data, calculating the silhouette only on the top five most common cell types. “*C. elegans*” shows the average silhouette values of the *C. elegans* scATAC-seq data using the broad cell types. “*C. elegans* neurons” shows the average silhouette value of only the neurons using the broad cell types. “*C. elegans* neuron subtypes” shows the average silhouette values using the detailed neuronal subtypes. “C. elegans neuron subtypes positive” shows the average silhouette values for the detailed neuronal subtypes, but averaging only across positive values.

We also attempted to transfer information across single cell data modalities by deriving a matrix prior from scRNA-seq data and applying it to a target scATAC-seq dataset. We found that with moderate values of the concentration parameter, the agreement between the outputs of LDA based on the target dataset and LDA based on the full data set improved when using the matrix prior compared to the uniform prior (Figure S12). Leveraging information from multiple single cell data modalities is an area of active research. Some popular tools, like Seurat [Butler et al., 2018], scVI Tools [Gayoso et al., 2021], or LIGER [Liu et al., 2020], take multiple data sets as input and use embedding techniques to analyze them jointly. These approaches are powerful, but require manipulating potentially very large datasets every time one wants to add new data into the model. In contrast, our matrix prior approach requires just a single large upfront compute task to train an LDA on a reference dataset, which yields a compact gene-topic matrix that can be used as a matrix prior for training comparatively lightweight LDA models in all subsequent analyses of new datasets. We also anticipate that new approaches, such as Polarbear [Zhang et al., 2022] and BABEL [Wu et al., 2021], that use deep learning models to translate data from one single cell modality to another will improve our ability to generate cross-modality matrix priors, not only between scATAC-seq and scRNA-seq data, but also between other pairs of modalities.

