## Supplementary Material for "Matrix prior for data transfer between single cell data types in latent Dirichlet allocation"

### Supplementary note 1: SHARE-seq scRNA-seq matrix prior may have limited use

We investigated the feasibility of using scRNA-seq data to construct a matrix prior for scATAC-seq analysis. This required that we map the scRNA-seq data and the scATAC-seq data onto a shared feature axis. To accomplish this mapping, translated the scATAC-seq data from a vocabulary of peaks to one of genes by simply counting the number of scATAC-seq cut sites overlapping each gene and its promoter (see Methods). We then proceeded with a similar analysis to Sections 4.2 and 4.3. To evaluate the performance of the matrix prior, we leveraged the fact that the scATAC-seq data and scRNA-seq data were generated from the same set of cells, and compared the inferred cell-topic and topic-gene matrices from the matrix prior LDA that was trained on the target scATAC-seq dataset to the output matrices generated by a joint model on the full scRNA-seq dataset.

We found that using the scRNA-seq data to construct the matrix prior did not consistently improve inference of the cell-topic and topic-gene matrices, although we saw some improvement at moderate values of $c_B$. For the cell-topic matrices, we found that as the concentration parameter $c_B$ increased beyond 4,000, the Pearson correlation to the joint model output tended to decrease. Similarly, Spearman correlation and MSE both got worse as $c_B$ increased beyond 4,000. For the topic-gene matrices, however, Spearman correlation improved as $c_B$ increased (Figures 2, S12). When plotting the raw values of the cell-topic matrix, we observed more points along the x- and y-axes. The reference model and joint model assigned low probability topics to different cells more frequently as $c_B$ increased (Figure S14). For the topic-gene matrix, the diagonal had increased density, meaning that the two analyses agreed more as $c_B$ increased (Figure S15)

We note that the read counts per gene and per cell in scRNA-seq was only moderately correlated with the number of cut sites summed over the gene body in the scATAC-seq data, with a Pearson correlation of 0.661 for the signal per cell, and 0.765 for the signal per gene (Figure S6). This makes sense because, although chromatin accessibility and gene expression are highly related biological phenomena, the information encoded by scATAC-seq and scRNA-seq count data is not the same. This could be one reason why the matrix prior only improves results compared to the uniform prior at moderate values of $c_B$. If a prior derived from a different data modality is given too much weight, then the discrepancies between the modalities are more likely to lead to a poor model fit.

### Supplementary Note 2: *C. elegans* silhouette values did not improve through use of the prior

The *C. elegans* data was previously labeled with a cell type based on marker genes through a clustering method that took into account all scATAC-seq peaks using all the cells together. We hypothesized that we would be better able to recover these cell type labels in the target dataset by incorporating information from the matrix prior. To this end, we analyzed the target dataset using both the matrix prior derived from a reference subset of cells and the uniform prior, and then evaluated how well the clusters in the cell-topic output agreed with the published cell type labels.

We used UMAP [McInnes et al., 2018] to reduce the 15-dimensional topic space to a two-dimensional representation and then colored each cell according to its cell type label. We compared these UMAP plots of the results of our LDA analysis with the matrix prior and with the uniform prior, as well as with different weights of $c_B$ (Figure S16a). The UMAP plots show that all of the LDA models produce reasonable agreement with the cell type labels, although we note that quantifying the cell type discrimination in each LDA output with the silhouette score shows a slight increase in the mean silhouette value, from 0.216 with the uniform prior to 0.220 with the matrix prior and $c_B = 4000$ (Figures 4, S16b).

Despite the similarity of the UMAP plots, we noted that at $c_B = 4000$ the neurons appeared to split into two clusters, and the average silhouette score for the neurons decreased from 0.254 with the uniform prior to 0.109 with the matrix prior and $c_B = 4000$. This observation suggested the hypothesis that the use of the prior might help to resolve fine grained cell types, so we repeated our cell type label analysis specifically on the neurons, this time using the published neuron subtype labels (Figure S17a). The UMAP plots show that as $c_B$ increased the cells were clustered more tightly together, however there was little change in how well the cell types were separated compared to the LDA with the uniform prior. In addition, the mean silhouette scores for the neuron subtypes (Figure S17b), did not increase as the value of $c_B$ increased. Overall, these analyses do not suggest that the neuron subtypes were further resolved by the use of the matrix prior.

| Variable | Definition |
|---|---|
| $N$ | The number of reads in a cell |
| $T$ | The number of topics |
| $V$ | The number of peaks or genes in the vocabulary |
| $U$ | The number of cells |
| $\boldsymbol{w}$ | A vector of genes or peaks in a cell |
| $\boldsymbol{\theta}$ | A vector $(\theta_1, \ldots, \theta_T)$; the topic distribution of a given cell |
| $\hat{\boldsymbol{\theta}}$ | Inferred $\boldsymbol{\theta}$ |
| $\boldsymbol{\phi}$ | A matrix such that $\phi_{tw}$ is the probability of observing a peak or gene $w$ for a topic $t$ |
| $\hat{\boldsymbol{\phi}}$ | Inferred $\boldsymbol{\phi}$ |
| $\boldsymbol{\alpha}$ | A basis vector of length $T$ that parameterizes the Dirichlet prior distribution over the cell-topic matrix. |
| $\boldsymbol{\beta}$ | A basis vector of length $V$ that parameterizes the Dirichlet prior distribution over the topic-peak/gene matrix. |
| $c_\alpha$ | The concentration parameter for $\boldsymbol{\alpha}$ |
| $c_\beta$ | The concentration parameter for $\boldsymbol{\beta}$ |
| $\boldsymbol{z}$ | A vector of topic assignments corresponding to $\boldsymbol{w}$ |
| $\xi$ | The average number of peaks or genes in a cell |
| $\boldsymbol{B}$ | A $T \times V$ matrix that parameterizes the matrix prior for the topic-peak/gene matrix. |
| $c_B$ | A concentration parameter for $\boldsymbol{B}$ |
| $\hat{\boldsymbol{\phi}}_{ref}$ | The output inferred $\boldsymbol{\phi}$ of an LDA analysis on the reference dataset. |

Table S1: Glossary of variables used

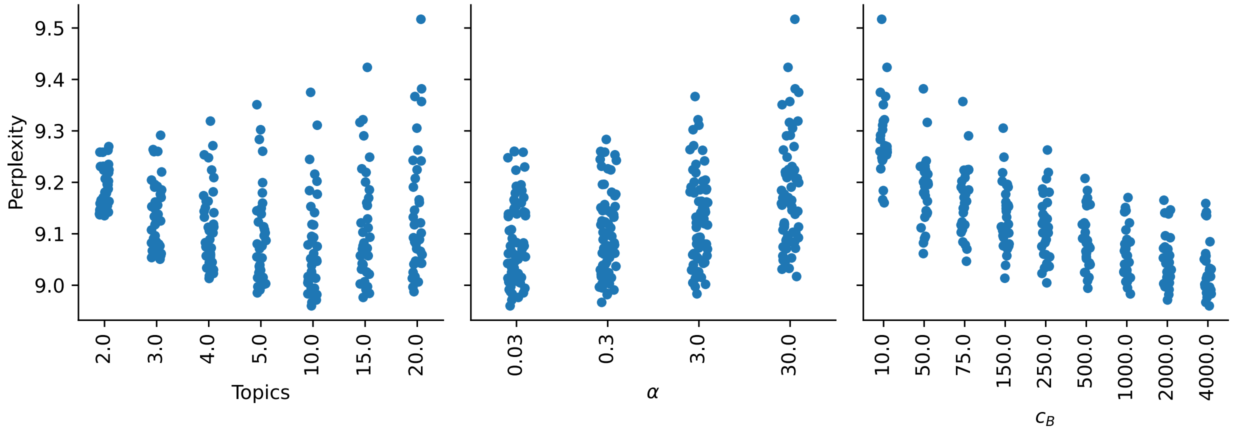


Figure S1: Hyperparameter search for *C. elegans* data was used to optimize the parameters. Each point is the average of 10 folds of the perplexity value of the test set. The x-axis is the value of the hyperparameter, and the y-axis is the perplexity value.

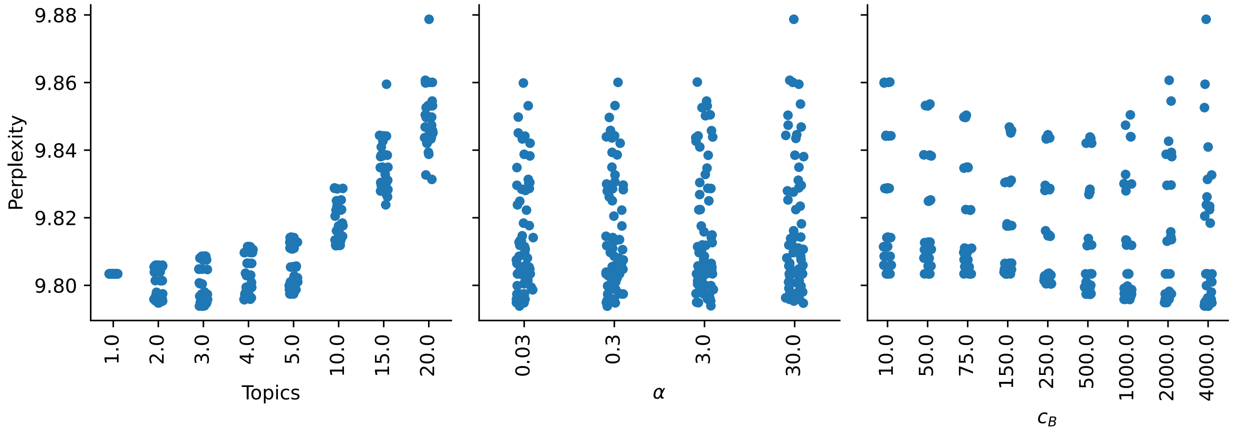


Figure S2: Hyperparameter search for SHARE-seq data with peaks was used to optimize the parameters. Each point is the average of 10 folds of the perplexity value of the test set. The x-axis is the value of the hyperparameter, and the y-axis is the perplexity value.

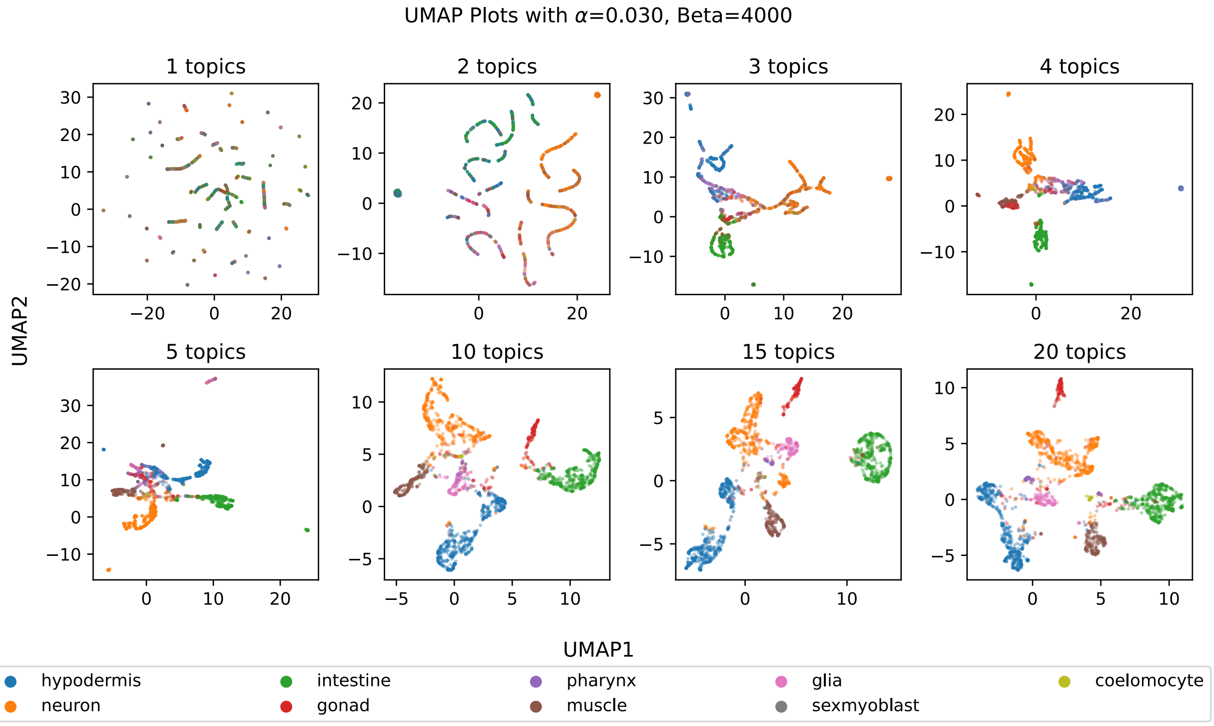


Figure S3: UMAP embeddings for different numbers of topics are shown to provide intuition on the effect of the number of topics. Embeddings were made for cell-topic matrices from matrix prior LDA on *C. elegans* scATAC-seq data using a fixed value of $c_B = 4,000$ and 13,734 peaks, while varying the number of topics.

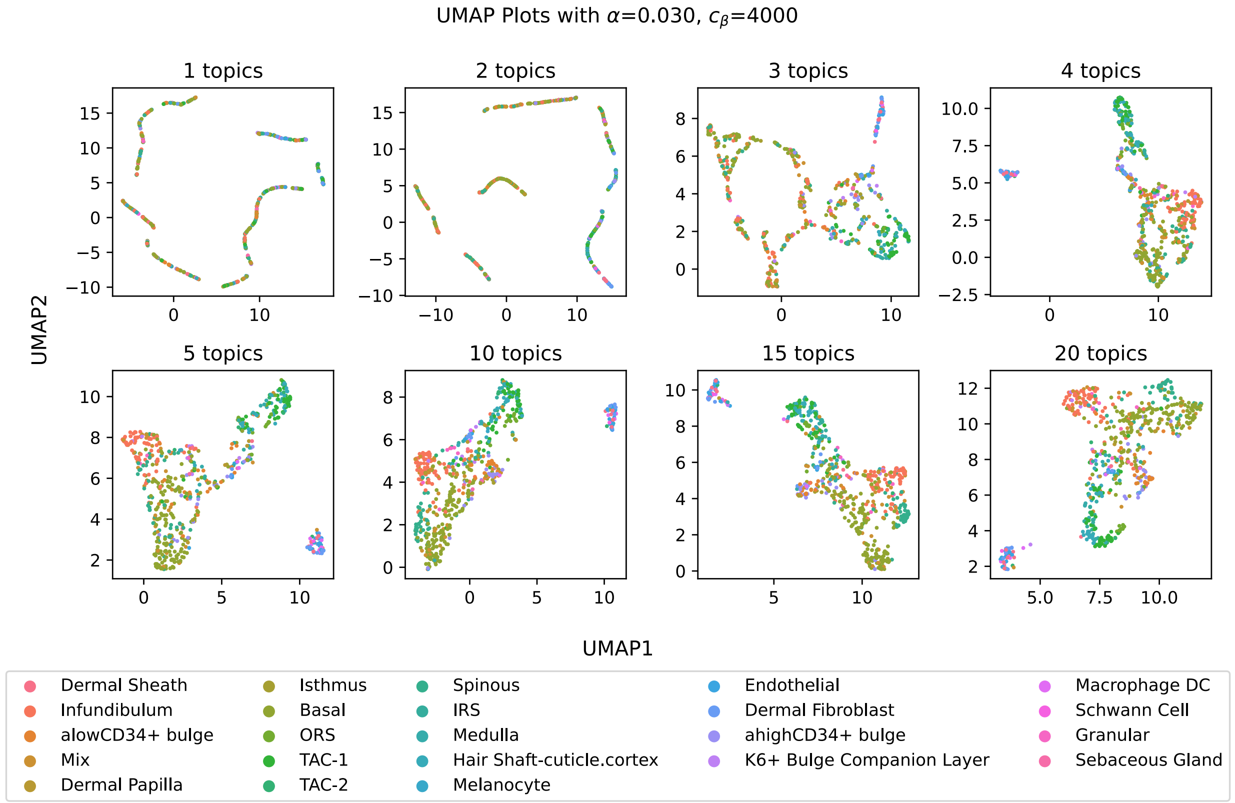


Figure S4: UMAP embeddings for different numbers of topics are shown to provide intuition on the effect of the number of topics. Plots were made with 630 target cells from the mouse skin SHARE-seq peak data subsetted to 7,000 cells and 20,000 most variable peaks. LDA was run with $c_\beta = 4,000$ using the uniform prior, while varying the number of topics.

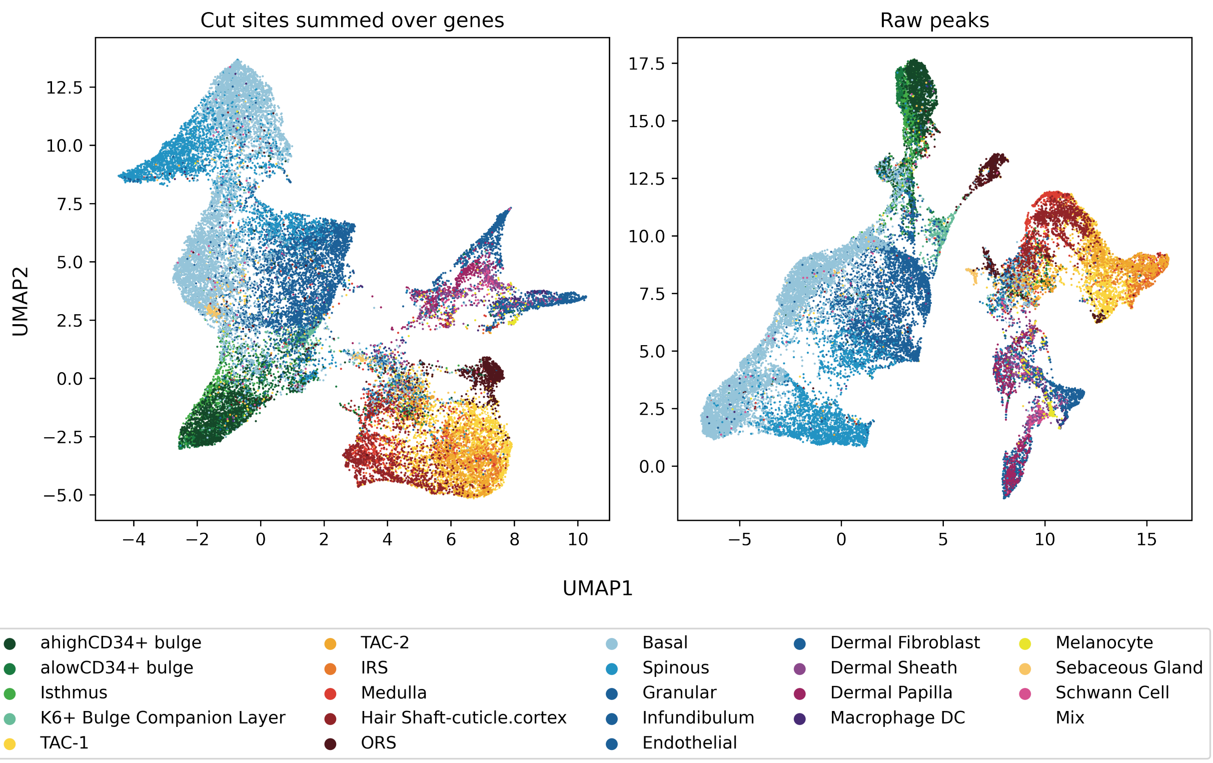


Figure S5: UMAP was applied to the cell-topic matrices output from LDA joint model to qualitatively compare cut sites summed over genes versus peaks. LDA was run with 15 topics, $c_\alpha = 3$, and $c_\beta = 4000$. On the left, the raw data fed into LDA are the cut sites summed over the 22,813 genes, as described in Section 3.2.4. On the right, the data fed into LDA are the 344,592 raw peaks.

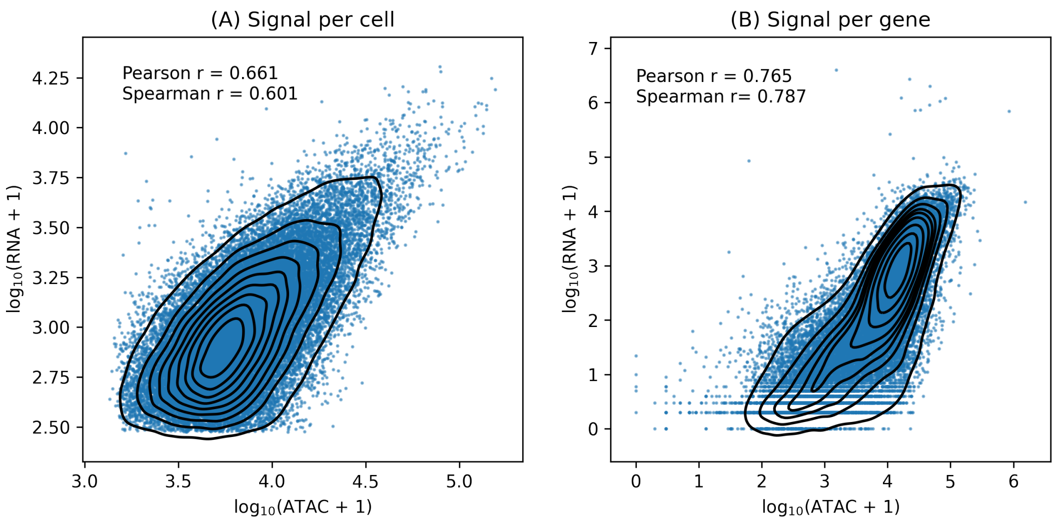


Figure S6: RNA reads and scATAC-seq cut sites summed over gene bodies are compared to determine the shared information between the data modalities. On the left, the signal per cell is the log plus one of the total number of counts for each cell (i.e. summing across all the genes in a cell). On the right, the signal per gene is the log plus one of the total number of counts for each gene (i.e. summing across all the cells for a gene). The Pearson correlation is reported in each plot. A kernel density estimator is overlayed on the data. The x-axis shows the score for the scATAC-seq data, and the y-axis shows the score for the scRNA-seq data.

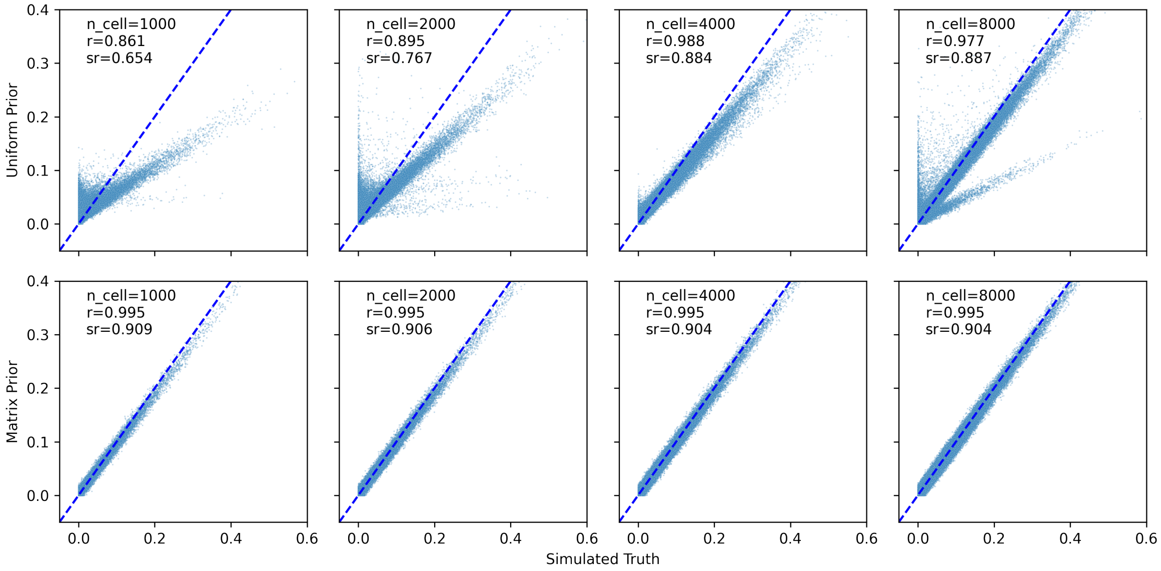


Figure S7: Scatter plots of cell-topic matrix values demonstrate the improvement of the matrix prior over the uniform prior in the true matrix simulation and further show that the performance of the uniform prior approaches that of the matrix prior as the number of cells increases. Plots show simulated true values (x-axis) of the cell-topic matrix against inferred values using LDA (y-axis). Pearson $r$ (r) and Spearman $r$ (sr) are reported for each plot. We compared different numbers of cells in the target dataset (different columns). We compared LDA with a uniform prior (top row) with a matrix prior generated from the true topic-gene matrix (bottom row). The blue dotted line is the line $y = x$.

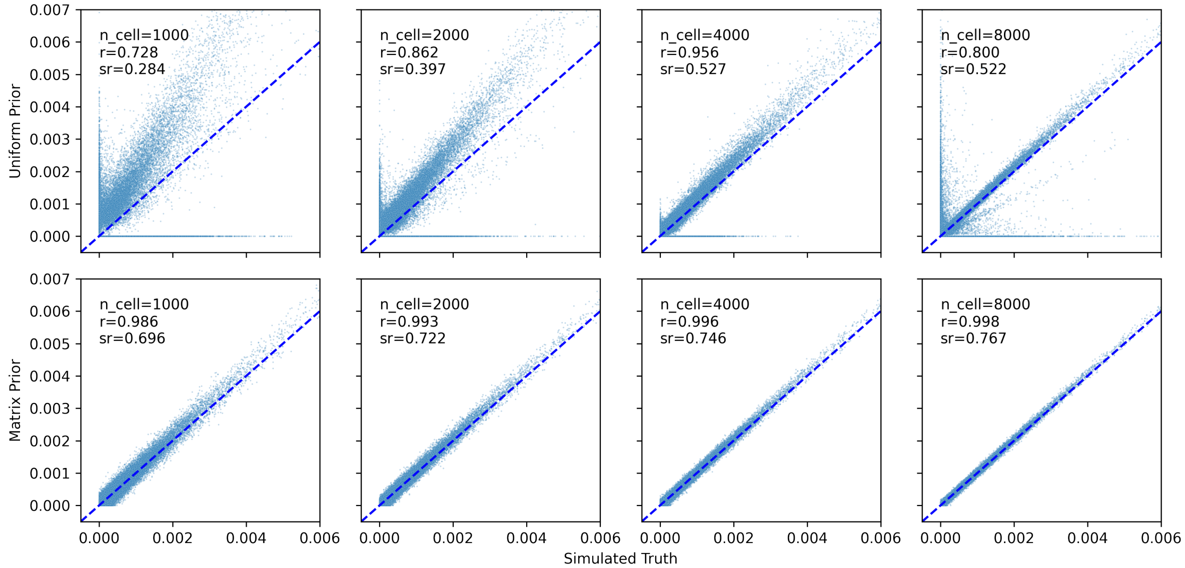


Figure S8: Scatter plots of topic-gene matrix values demonstrate the improvement of the matrix prior over the uniform prior in the true matrix simulation and further show that the performance of the uniform prior approaches that of the matrix prior as the number of cells increases Plots show simulated true values (x-axis) of the topic-gene matrix against inferred values using LDA (y-axis). Pearson $r$ (r) and Spearman $r$ (sr) are reported for each plot. We compared different numbers of cells in the target dataset (different columns). We compared LDA with a uniform prior (top row) with a matrix prior generated from the true topic-gene matrix (bottom row). The blue dotted line is the line $y = x$.

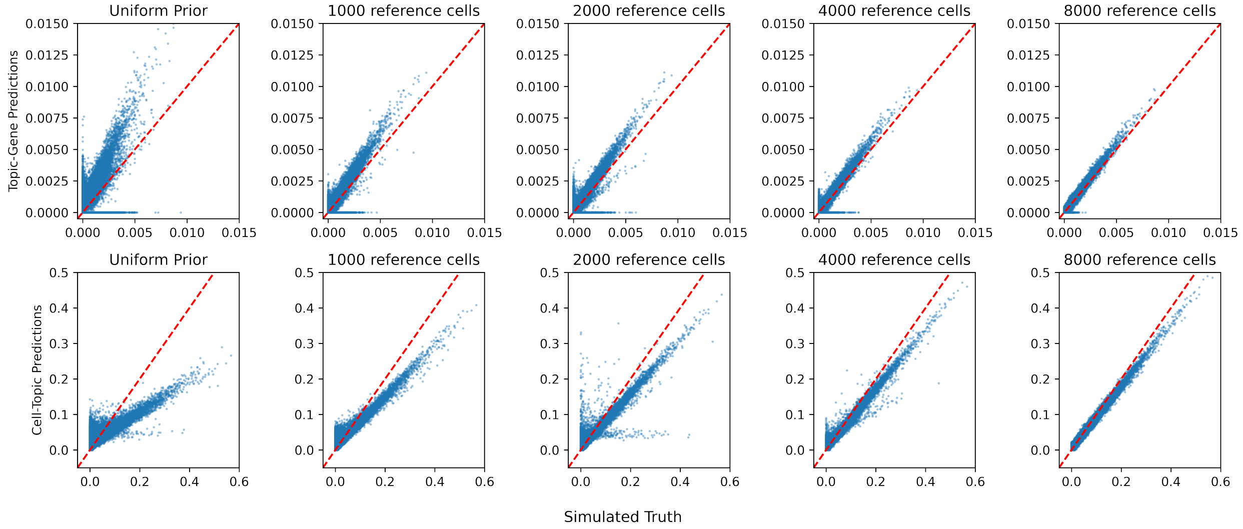


Figure S9: Scatter plots demonstrate the improvement of the matrix prior in both the cell-topic and topic-gene matrices as the number of reference cells increases in the inferred matrix simulation. 1000 simulated cells were analyzed using a uniform prior (left-most column) and a matrix prior. The dotted red line is the $y = x$ line. True simulated values (x-axis) and inferred values (y-axis) are plotted for both the topic-gene matrices (top) cell-topic matrices (bottom).

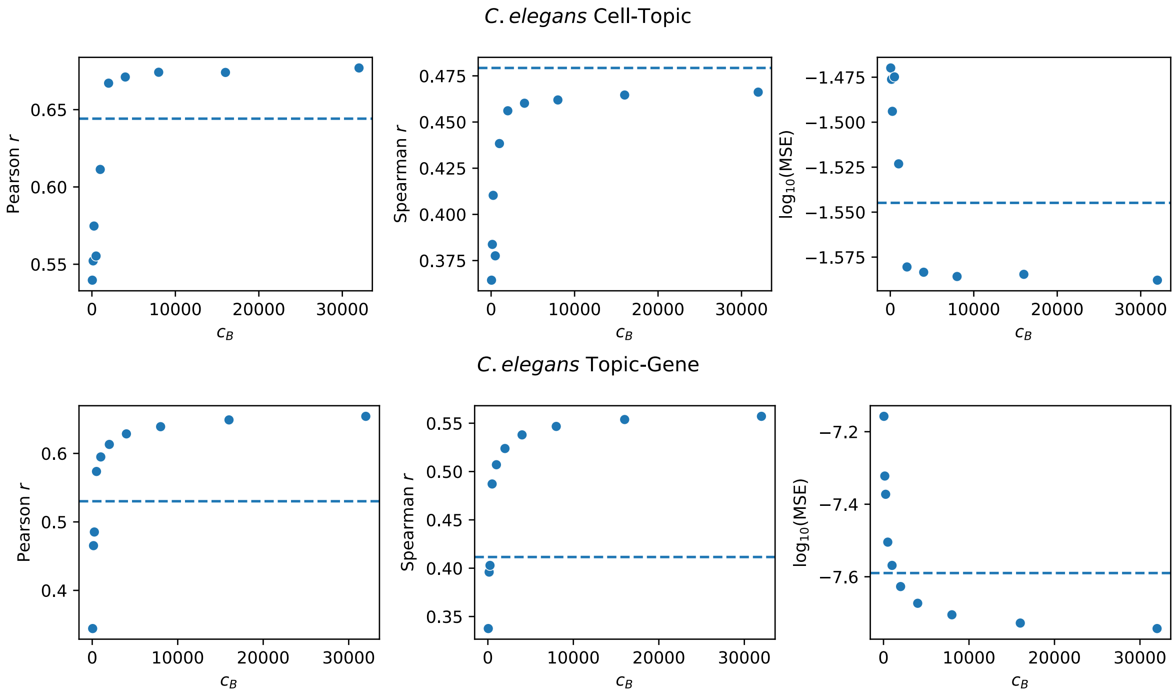


Figure S10: Summary statistics (y-axis) between output matrices of the joint model and LDA with matrix prior, both trained on the *C. elegans* data, show that as the weight of the prior increases, agreement between the matrix prior and joint model also increases. Each plot shows the matrix prior LDA results (points) for increasing values of $c_B$ (x-axis) versus the uniform prior (blue dotted line). The top row of plots shows summary statistics for the cell-topic matrix, and the bottom row of plots shows summary statistics for the topic-gene matrix.

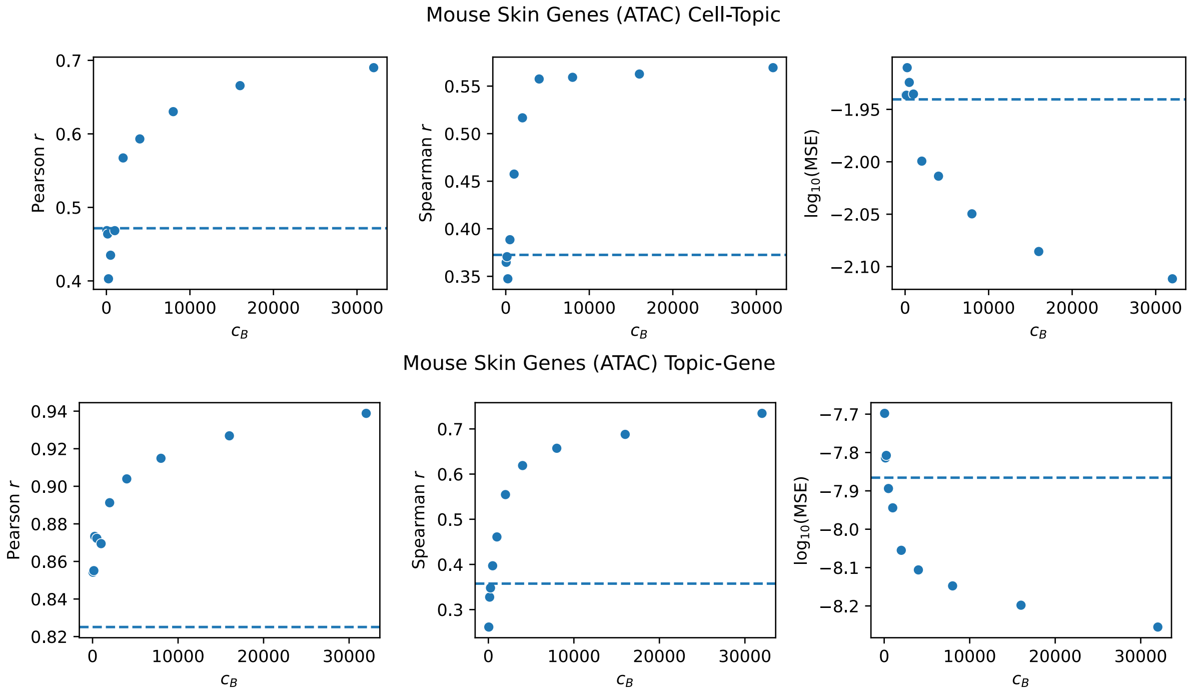


Figure S11: Summary statistics (y-axis) between output matrices of the joint model and LDA with matrix prior, both trained on the SHARE-seq mouse skin scATAC-seq data with cut sites summed over genes (i.e. using the genes vocabulary), show that as the weight of the prior increases, agreement between the matrix prior and joint model also increases. Each plot shows the matrix prior LDA results (points) for increasing values of $c_B$ (x-axis) versus the uniform prior (blue dotted line). The top row of plots shows summary statistics for the cell-topic matrix, and the bottom row of plots shows summary statistics for the topic-gene matrix.

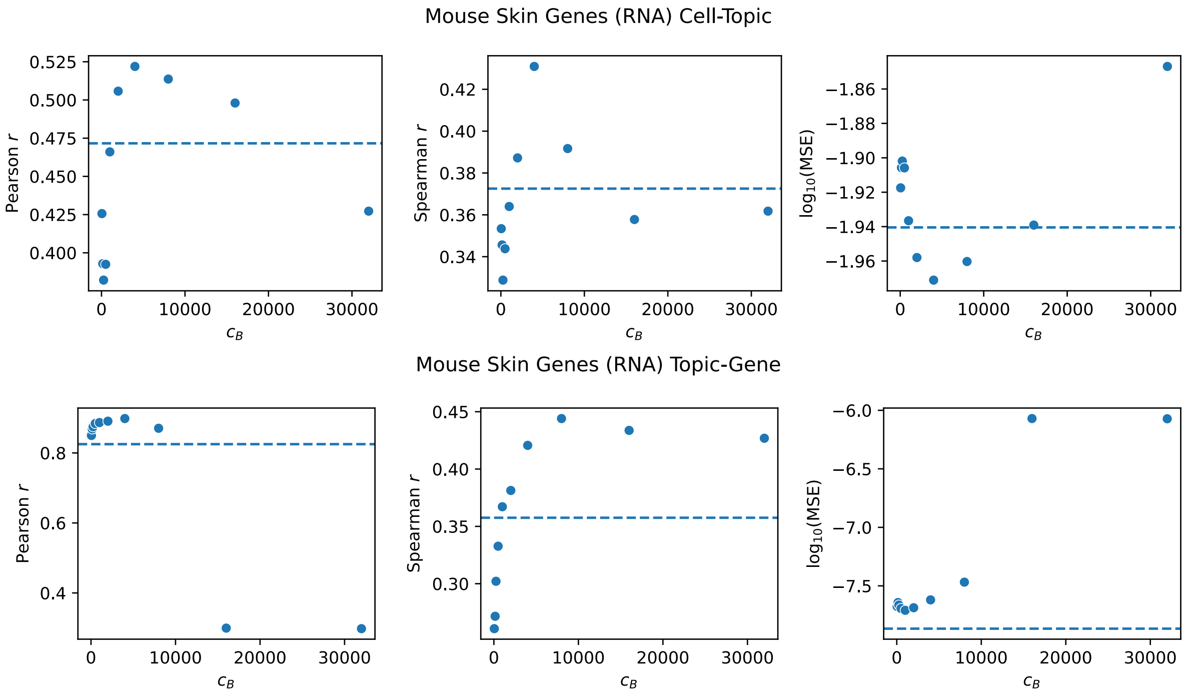


Figure S12: Summary statistics (y-axis) between output matrices of the joint model and LDA with matrix prior, both trained on the SHARE-seq mouse skin scRNA-seq data, show that as the weight of the prior increases, agreement between the matrix prior and joint model also increases with moderate weights and declines with higher weights. Each plot shows the matrix prior LDA results (points) for increasing values of $c_B$ (x-axis) versus the uniform prior (blue dotted line). The top row of plots shows summary statistics for the cell-topic matrix, and the bottom row of plots shows summary statistics for the topic-gene matrix.

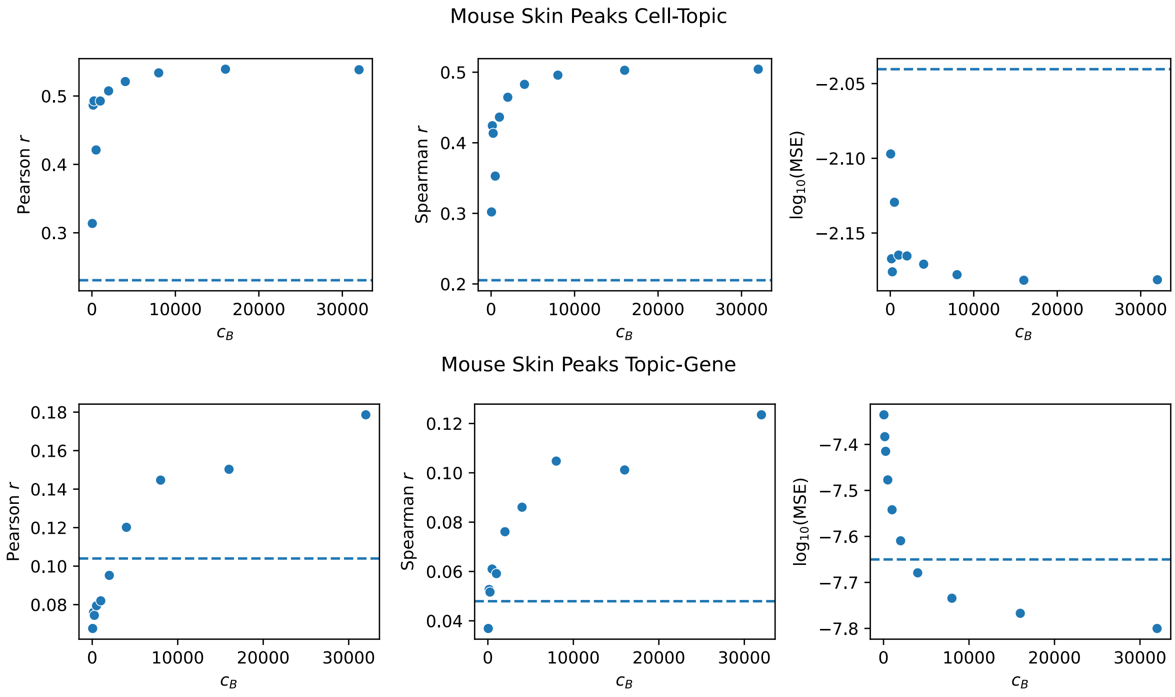


Figure S13: Summary statistics (y-axis) between output matrices of the joint model and LDA with matrix prior, both trained on the SHARE-seq mouse skin scATAC-seq data (i.e. using the peaks vocabulary), show that as the weight of the prior increases, agreement between the matrix prior and joint model also increases. Each plot shows the matrix prior LDA results (points) for increasing values of $c_B$ (x-axis) versus the uniform prior (blue dotted line). The top row of plots shows summary statistics for the cell-topic matrix, and the bottom row of plots shows summary statistics for the topic-gene matrix.

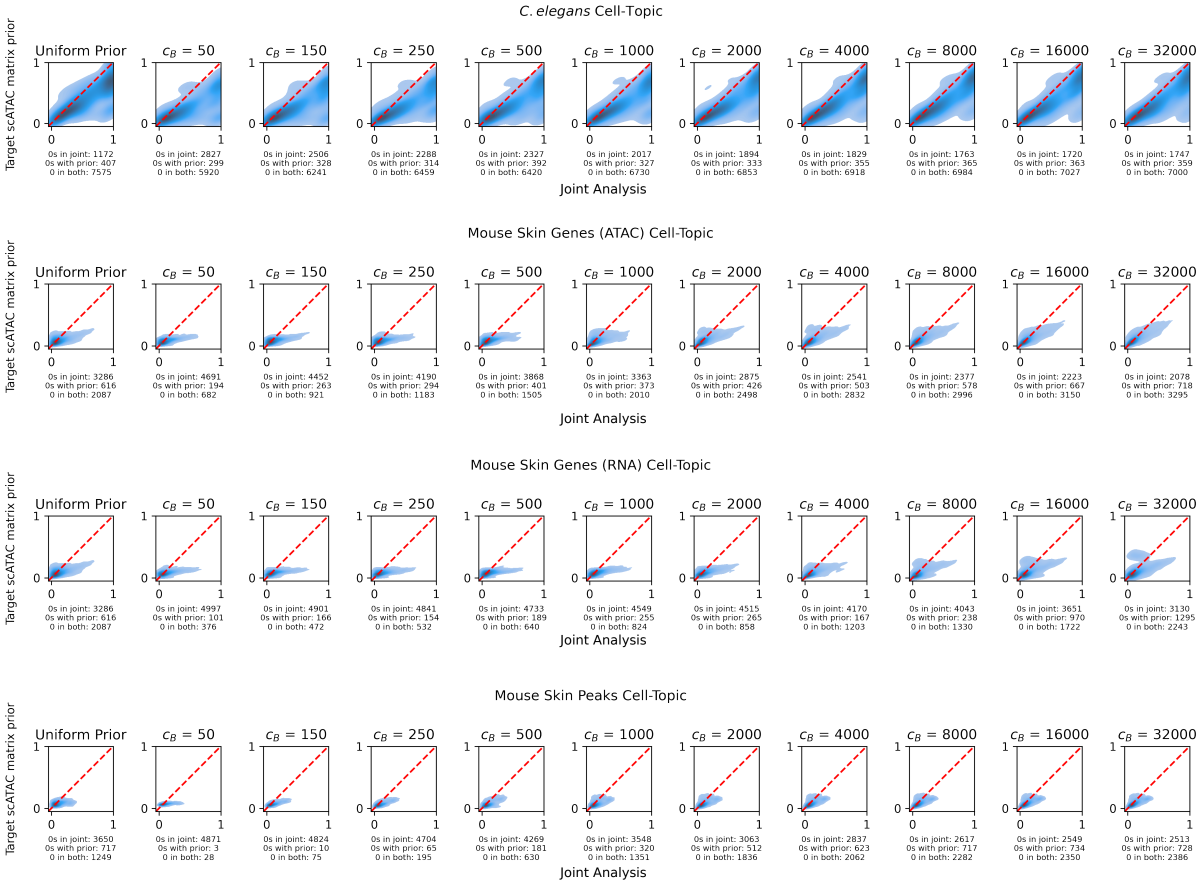


Figure S14: Comparing the cell-topic matrices of the joint model versus the LDA with the matrix prior reveals that as the weight of the matrix prior increases, the agreement between the models increases. The effect of different values of $c_B$ were evaluated by comparing the cell-topic matrix using the matrix prior to the cell-topic matrix from the joint model. Different values of $c_B$ are plotted across different columns, and different datasets are shown in different rows. We first flatten the cell-topic matrices so that they can be plotted. The cell-topic assignments from the joint model are shown on the x-axis, and the inferred cell-topic assignments from the matrix prior LDA are shown on the y-axis. A dotted red line is drawn to indicate the line $y = x$. Zero values are omitted from the plots, but the number of zeros exclusively in the cell-topic matrix of the joint model, exclusively in the cell-topic matrix of the LDA with matrix prior, and number of zeros in both is noted below each plot.

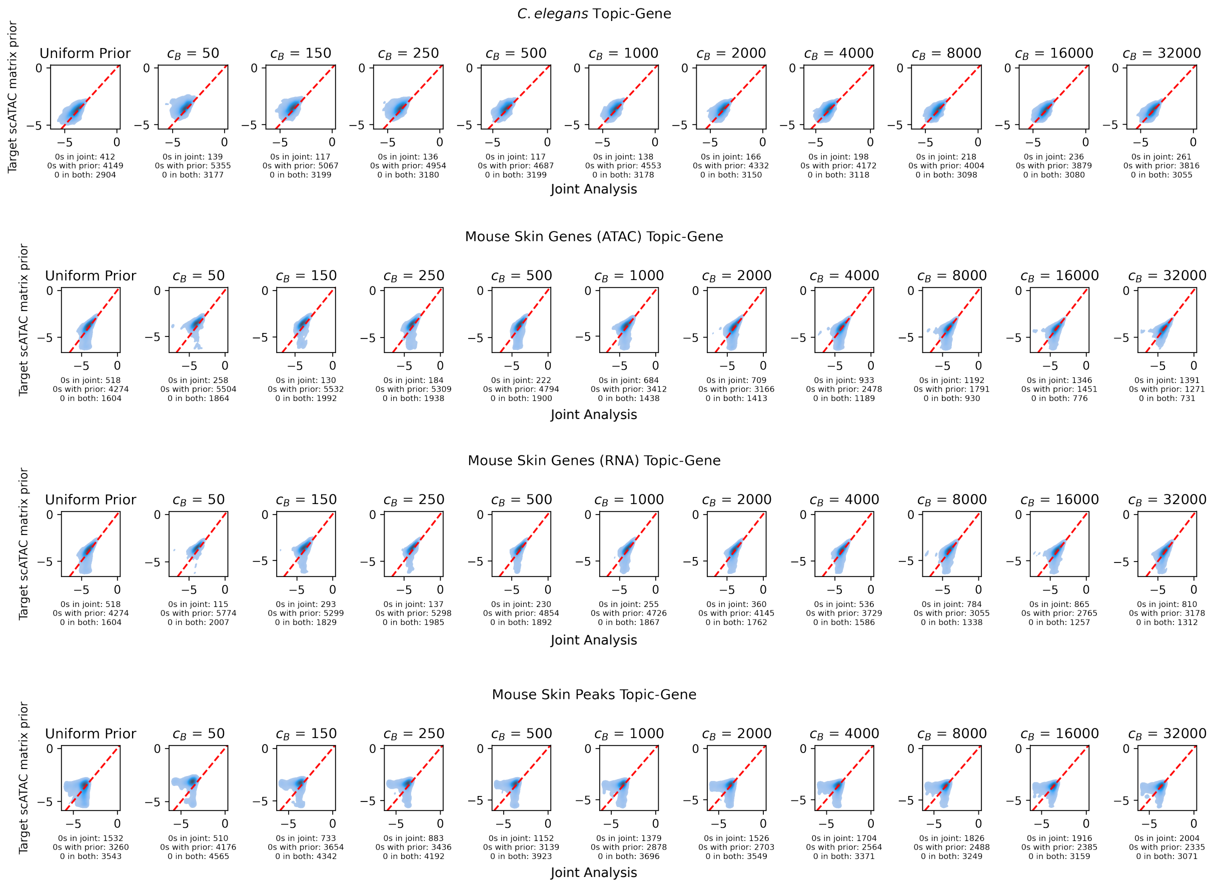


Figure S15: Comparing the topic-gene matrices of the joint model versus the LDA with the matrix prior reveals that as the weight of the prior increases, the agreement between the models increases. The effect of different values of $c_B$ were evaluated by comparing the topic-gene matrix using the matrix prior to the topic-gene matrix from the joint model. Different values of $c_B$ are plotted across different columns, and different datasets are shown in different rows. We first flatten the topic-gene matrices so that they can be plotted. The topic-gene assignments from the joint model are shown on the x-axis, and the inferred topic-gene assignments from the matrix prior LDA are shown on the y-axis. A dotted red line is drawn to indicate the line $y = x$. Zero values are omitted from the plots, but the number of zeros exclusively in the topic-gene matrix of the joint model, exclusively in the topic-gene matrix of the LDA with matrix prior, and number of zeros in both is noted below each plot.

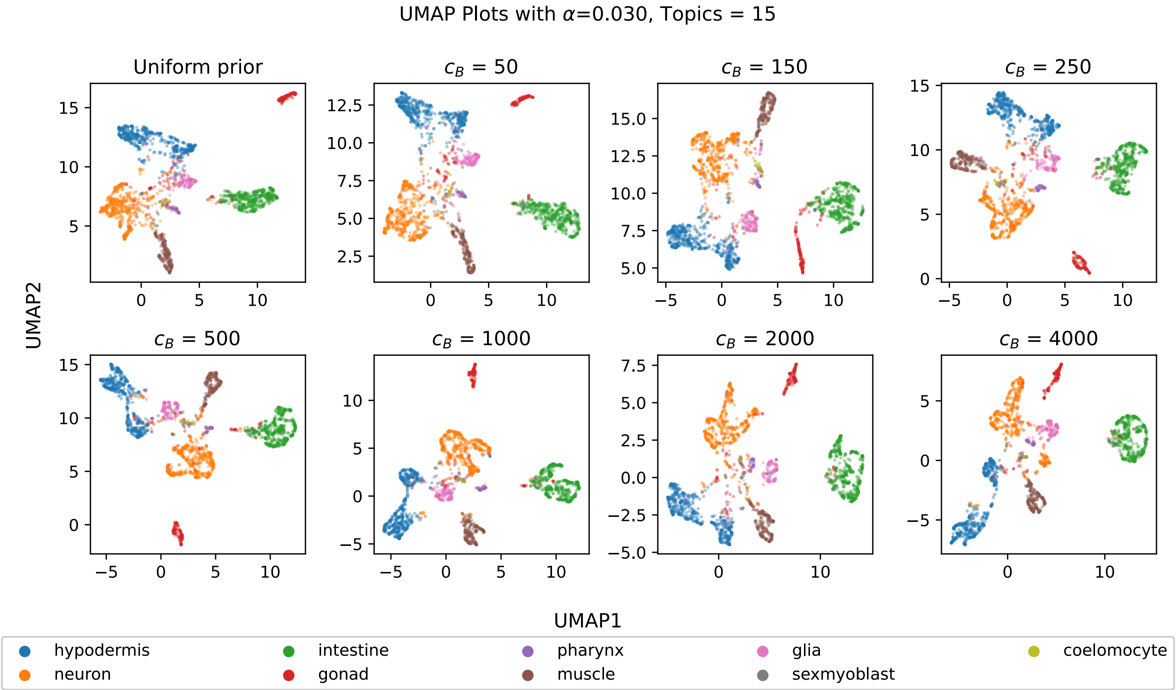


(a) UMAP plot of all the *C. elegans* data reveal cell type structure from LDA analysis with different weights $c_B$ of the matrix prior. Cells are colored based on their published cell type annotations.

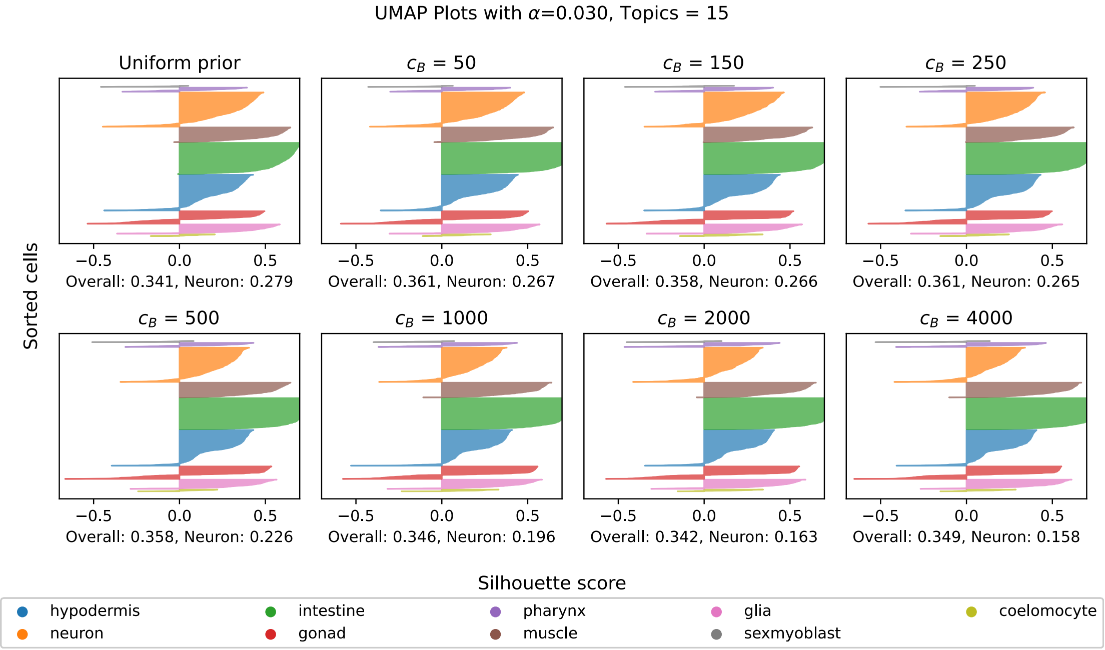


(b) Silhouette plots demonstrate that in *C. elegans*, the silhouette value did not improve with increased weight of the prior. Silhouette values are shown for *C. elegans* cell types plotted for results from a uniform prior LDA model and matrix prior LDA models trained with increasing values of $c_B$ using 15 topics and the scATAC-seq data translated into the genes vocabulary (ATAC cut sites summed over the promoter and gene body for 13,734 genes). Each row in each plot represents one cell, and the silhouette value of the cell is the length of the line. The mean silhouette value for all of the cells is shown as "Overall", and the mean silhouette value for only the neurons is shown as "Neuron."

Figure S16: UMAP of the cell-topic matrices (a) and silhouette plots (b) show that increased weight of the matrix prior does not seem to improve the ability of the matrix prior LDA to distinguish among cell types in the *C. elegans* scATAC-seq data.

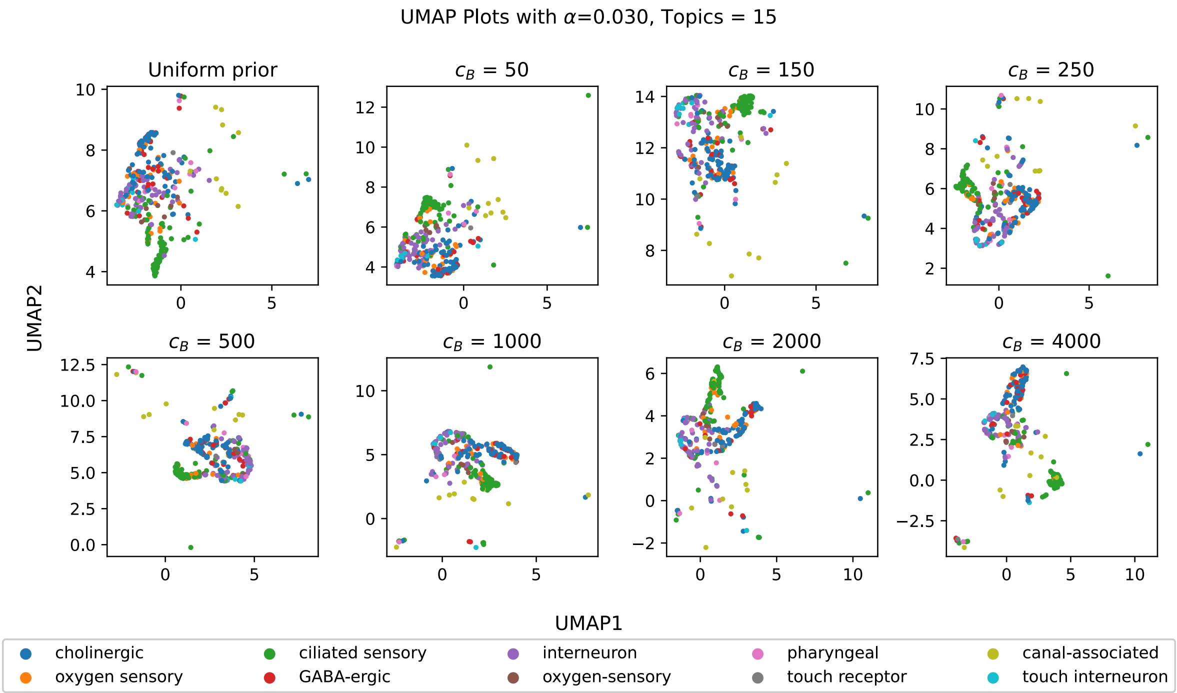


(a) A subset of Figure S16a that includes only neurons, demonstrating that increased values of $c_B$ have little effect on the ability of the matrix prior LDA to distinguish among published cell types. Cells are colored by published neuron subtype labels.

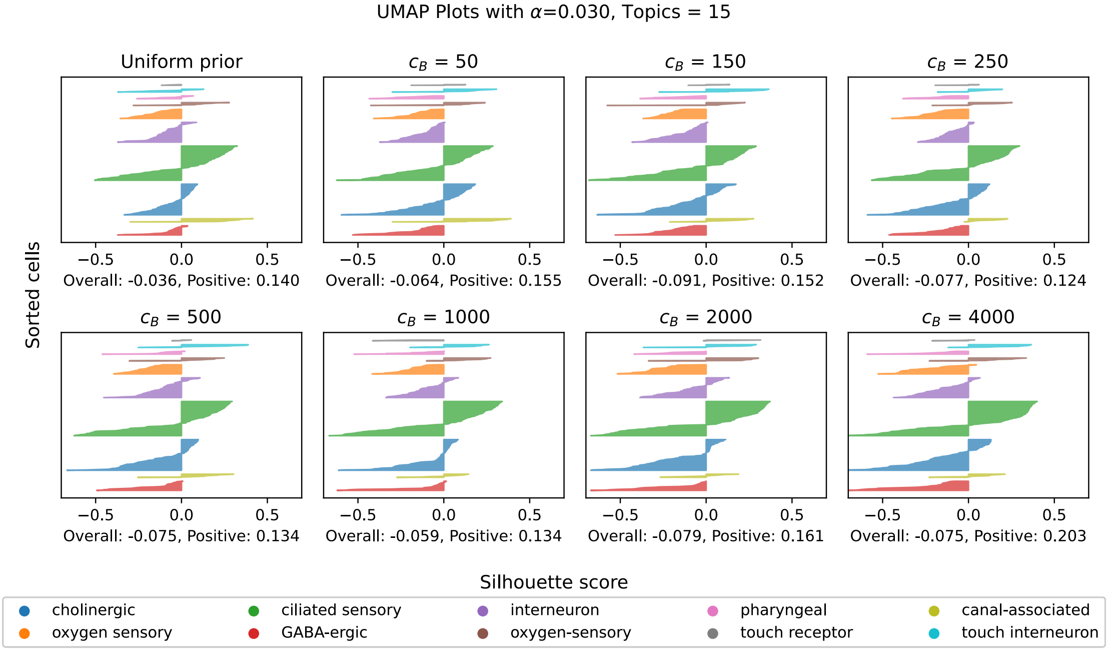


(b) Silhouette plots of the neurons in the *C. elegans* dataset. The overall mean silhouette values and the mean positive silhouette values are reported.

Figure S17: UMAP of the cell-topic matrices (a) and silhouette plots (b) show that increased weight of the matrix prior does not seem to improve the ability of the matrix prior LDA to distinguish among fine grained cell types in the neurons from the *C. elegans* scATAC-seq data.

| Cell Type | Count |
|---|---|
| Basal | 159 |
| Infundibulum | 74 |
| TAC-1 | 56 |
| Spinous | 52 |
| Mix | 46 |
| alowCD34+ bulge | 32 |
| Hair Shaft-cuticle.cortex | 29 |
| Endothelial | 22 |
| ahighCD34+ bulge | 22 |
| Medulla | 19 |
| Dermal Fibroblast | 18 |
| ORS | 18 |
| Isthmus | 13 |
| IRS | 12 |
| Dermal Sheath | 11 |
| TAC-2 | 10 |
| Dermal Papilla | 8 |
| Macrophage DC | 8 |
| K6+ Bulge Companion Layer | 7 |
| Granular | 6 |
| Schwann Cell | 3 |
| Melanocyte | 3 |
| Sebaceous Gland | 2 |

Table S1: Table of counts of mouse skin cell types used in the target dataset.

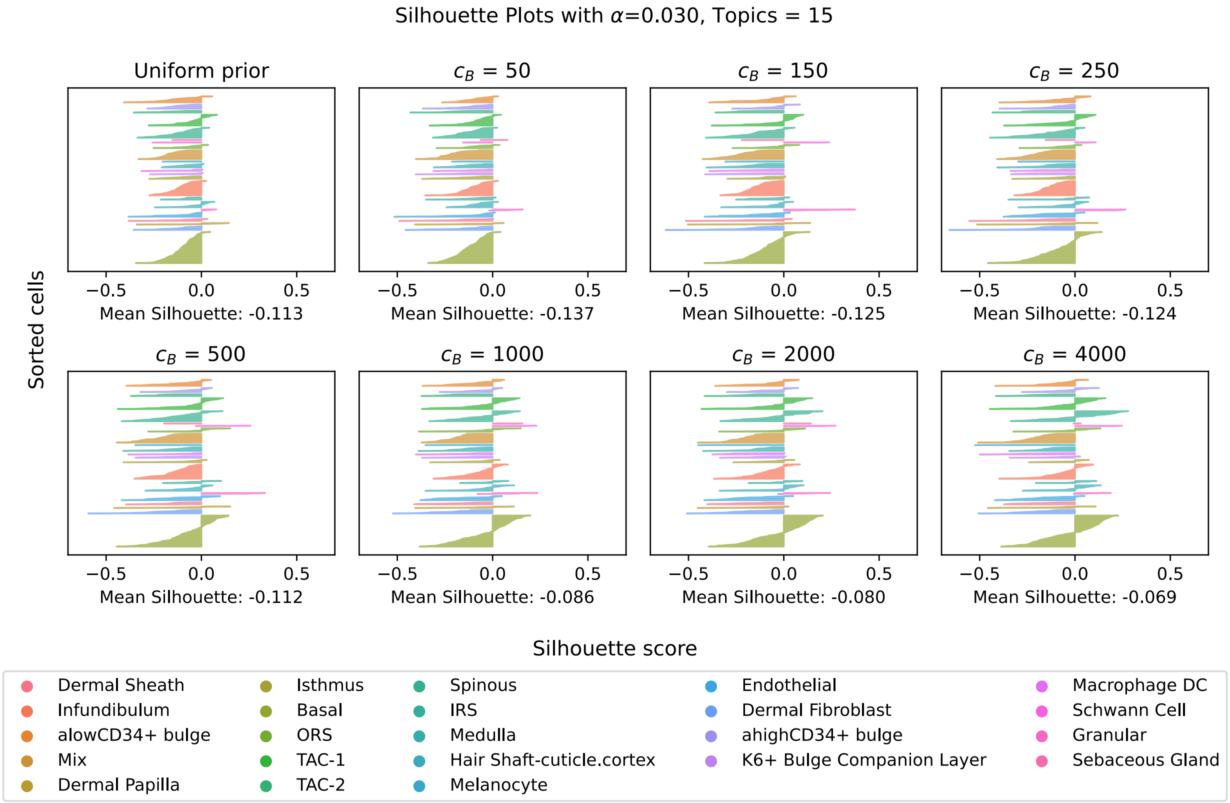


Figure S18: Silhouette scores calculated from matrix prior LDA models of the SHARE-seq mouse skin scATAC-seq data demonstrate better separation of cell type clusters with higher weights for the matrix prior ($c_B$). Plots showing silhouette values for results from a uniform LDA model and matrix prior LDA models with increasing values of $c_B$, all trained with 15 topics and the randomly-selected 20000 scATAC-seq peaks.

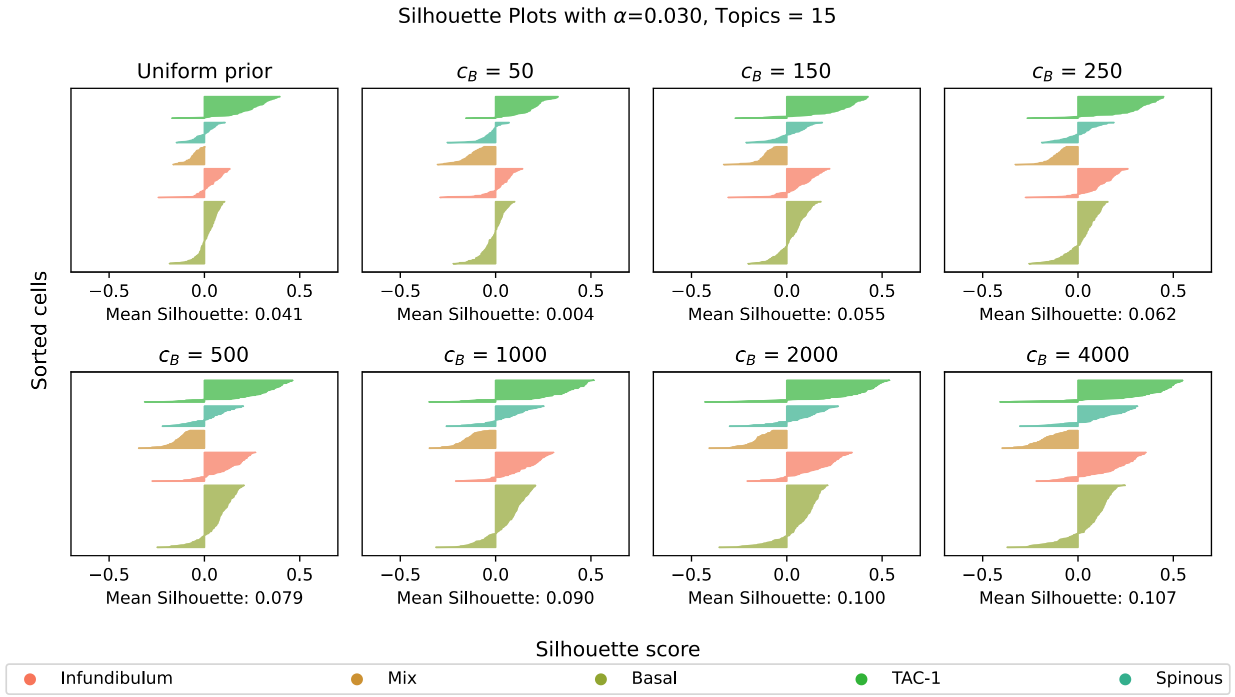


Figure S19: Silhouette scores calculated for just the top 5 most common mouse skin cell types based on the same models in Figure S18 show positive mean silhouette score values that increase as $c_B$ increases, suggesting higher matrix prior weights improve the ability of the matrix prior LDA to distinguish among cell types.
